# Protective effects of Mkl1/2 against lipodystrophy and muscle atrophy via PI3K/AKT-independent FoxO repression

**DOI:** 10.1101/2023.08.09.552644

**Authors:** Atsushi Kubo, Keisuke Hitachi, Ryutaro Shirakawa, Toshihiko Ogura

## Abstract

FoxO transcription factors are involved in the pathogenesis of lipodystrophy and muscle atrophy. FoxO proteins are phosphorylated and inactivated by PI3K/AKT signaling; however, little is known about FoxO repressors other than this pathway. Our study showed that the Srf cofactors Mkl1 and Mkl2 directly repressed FoxO transcriptional activity, independent of the PI3K/AKT pathway. Loss of *Mkl1/2* led to the overactivation of FoxO, which impaired the maintenance of both white adipose tissue (WAT) and skeletal muscle. In *Mkl2*-deficient preadipocytes, *Pparγ* was suppressed and white adipogenesis was severely impaired. In myotubes, Mkl1 and Mkl2 suppressed the expression of atrophy-related genes (atrogenes) induced by FoxO. *Mkl1* expression was reduced at the onset of muscle atrophy *in vivo*, and exogenous supplementation of skeletal muscle Mkl1 suppressed atrogene expression and induced muscle hypertrophy. Finally, both Mkl2 nuclear localization and *Mkl1/2* expression were upregulated by exercise, suggesting that *Mkl1/2* is involved in the inhibitory effect of exercise on muscle atrophy. These findings indicate that Mkl1/2 act as PI3K/AKT signaling-independent repressors of FoxO and are essential for adipose and muscle tissue homeostasis.

## INTRODUCTION

FoxO proteins are involved in many aspects of life^1, 2^. In lower organisms, FoxO activation has been reported to increase the lifespan^3^, and in mammals, there is emerging evidence that excessive FoxO activation is likely to induce cellular senescence or age-related diseases^4, 5^. As FoxO proteins are known to be phosphorylated and inactivated by the PI3K/AKT pathway^6^, excessive FoxO activation is often associated with reduced responsiveness to insulin signaling. Adipose tissue and skeletal muscle are tissues closely associated with insulin resistance.

Excessive activation of Foxo1 in white adipose tissue (WAT) affects systemic metabolism and is implicated in the development of diabetes^5^. In addition, the subcutaneous fat mass decreases with age. Since subcutaneous fat is thought to have beneficial effects on systemic metabolism, some age-related metabolic dysfunctions may also be due to a lack of subcutaneous fat^7^. WAT maintains its plasticity through the differentiation of adipose stem and progenitor cells (ASPCs), but the differentiation capacity of aged ASPCs is reduced^7, 8^. Foxo1 overactivation reportedly negatively regulates both the initiation of *Pparγ* expression and the transcriptional activity of Pparγ itself, thereby suppressing adipocyte differentiation^9–12^.

In addition to *Foxo1*, *Foxo3* and *Foxo4* have overlapping functions in skeletal muscle. Skeletal muscle atrophy can be caused by a variety of factors, including aging, fasting, denervation, cancer cachexia, and glucocorticoid treatment, all of which are closely related to the activation of FoxO proteins^13–16^. Activated Foxo1/3/4 promotes proteasomal proteolysis of muscle proteins by inducing the expression of atrophy-related genes (atrogenes), including skeletal muscle-specific E3 ubiquitin ligases (*atrogin-1* and *MuRF1*)^16, 17^. Glucocorticoids and glucocorticoid receptor (GR) are also known to be involved in skeletal muscle atrophy. Glucocorticoids activate FoxO by dephosphorylation and increase its expression^13, 18, 19^. GR and FoxO synergistically induce atrogenes^20^.

In contrast, exercise induces muscle hypertrophy and is an effective intervention against muscle wasting^21^. Increased insulin responsiveness due to exercise is thought to influence muscle hypertrophy and atrophy suppression; however, the exact mechanisms are not well understood. Exercise upregulates the expression of *PGC-1α*^22, 23^, the overexpression of which protects against muscle wasting^24, 25^. However, overexpression of *PGC-1α* also increases Akt protein levels, resulting in increased Foxo3 phosphorylation^26^. Therefore, the inhibitory effect of *PGC-1α* overexpression on muscle atrophy may be mediated, in part, through the enhancement of the PI3K/AKT pathway.

*Mkl1* and *Mkl2* play important roles in the cardiovascular system by acting as cofactors for *Srf*^27, 28^. These are thought to have overlapping molecular functions. However, in contrast to *Mkl1*-knockout (KO) mice, *Mkl2*-KO mice show embryonic lethality^29, 30^. *Mkl1* and *Mkl2* are mechanosensitive transcription factors. Their subcellular localization (i.e. whether or not they are activated) is also regulated by intracellular actin dynamics^27, 28^. G-actin and importin competitively bind in the region around the RPEL domain of Mkl1 and Mkl2. Once actin polymerization occurs, such as that induced by mechanical stress, it leads to the nuclear accumulation of Mkl1/2 through a decrease in G-actin binding.

The association with laminopathy is another distinguishing feature of *Mkl1/2*. Mutations in *Lmna*, an intermediate filament protein that forms the nuclear lamina, are known to cause a variety of inherited diseases, including premature aging, lipodystrophy, muscular dystrophy, and dilated cardiomyopathy (DCM)^31, 32^. *Lmna* deficiency or the *Lmna* N195K mutation is closely associated with DCM pathogenesis, and nuclear translocation of Mkl1 is reported to be severely suppressed in cells carrying these mutations^33^. Interestingly, excessive FoxO activation has been observed in the myocardium of mice with DCM associated with *Lmna* deficiency, and this dysregulation of *FoxO* is thought to be involved in disease progression^34^. However, the mechanism underlying FoxO overactivation caused by *Lmna* deficiency is not well understood.

In the present study, we showed that the *Srf* cofactors Mkl1 and Mkl2 repress FoxO protein activity through direct protein-protein interactions and help maintain WAT and skeletal muscle mass.

## RESULTS

### *Mkl2*-KO mice show a decrease in both WAT and skeletal muscle mass

*Mkl2*-KO mice of the C57BL/6 strain show embryonic lethality due to cardiovascular developmental abnormalities, including dysplasia of the sixth pharyngeal arch artery and ventricular septal defects^29, 30^. To investigate the function of *Mkl2* in adults, we generated *Mkl2*-KO mice by heterosis using crosses between ICR and C57BL/6N strains. *Mkl2*-KO mice weighed significantly less than wild-type mice in the same litter (Dunnett’s test, male mice: p=0.0023, female mice: p <0.0001) (Fig. 1a). Significant reductions in WAT were observed in the inguinal, epididymal, gonadal, and mesenteric regions (Fig. 1b, c, e, Extended Data Fig. 1a, c, d), whereas there was little change in the weight of brown adipose tissue (Extended Data Fig. 1b).

**Fig. 1:**
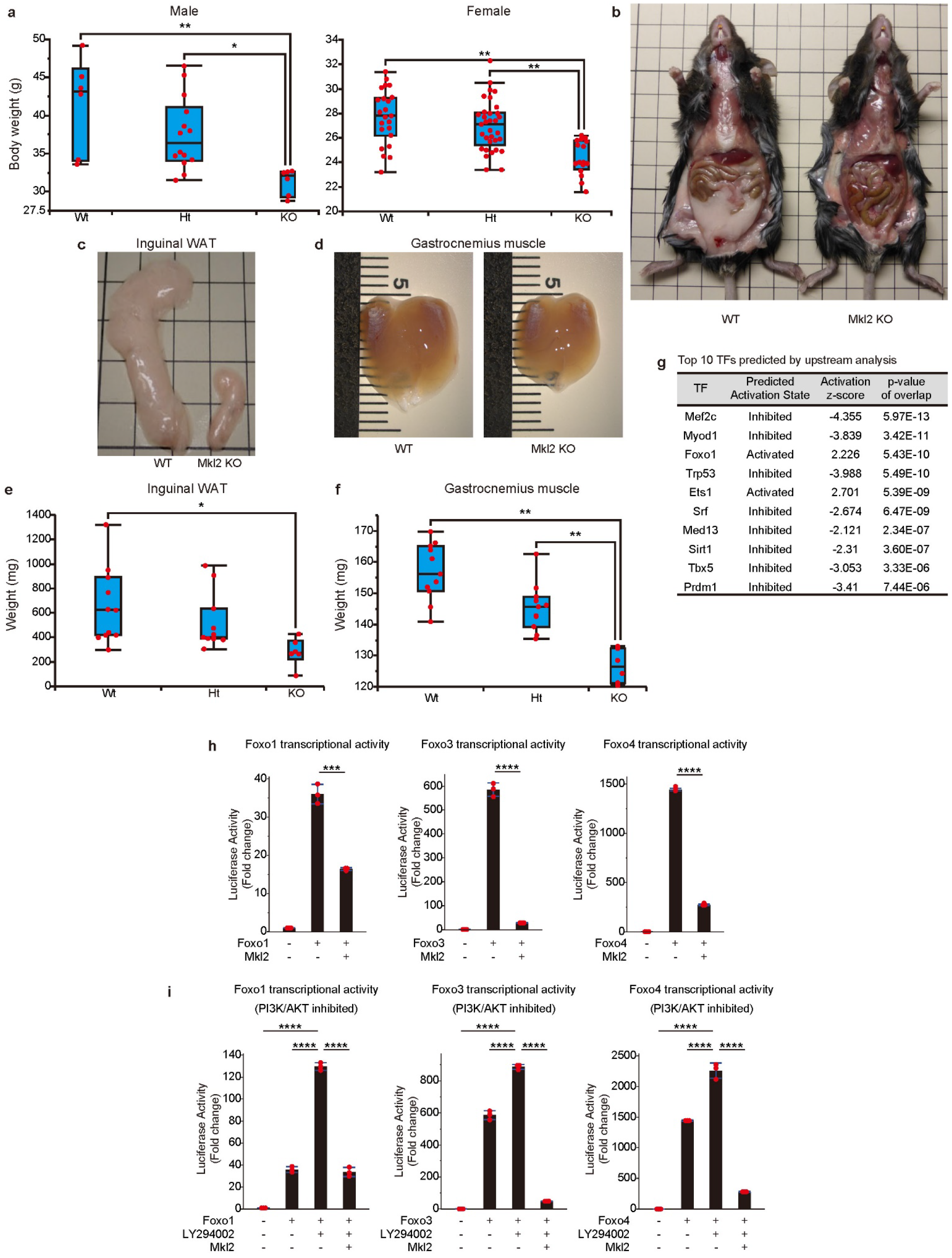
In *Mkl2*-knockout mice, both white adipose tissue and skeletal muscle mass are reduced. **a**, Body weight of *Mkl2* KO mice and their littermates. Data are presented as box plots (male: WT, n=6; Ht, n=17; KO, n=7; female: WT, n=23; Ht, n=33; KO, n=16). **b**, **c**, White adipose tissue (WAT) of *Mkl2* KO mice and their littermates. Whole abdomen (**b**) and inguinal WAT (**c**). **d**, Gastrocnemius muscle of *Mkl2* KO mice and their littermates. **e, f**, Tissue weights of *Mkl2* KO mice and their littermates. Inguinal WAT (**e**) and gastrocnemius muscle (**f**). Data are presented as box plots (21-22 weeks old, female mice; WT, n=11; Ht, n=11; KO, n=6). **g**, Prediction of upstream regulatory transcription factors based on differentially expressed genes in the inguinal WAT of *Mkl2* KO mice. **h**, **i**, Inhibitory effects of Mkl2 on the transcriptional activity of Foxo1, Foxo3, and Foxo4 using luciferase reporters with FoxO-responsive elements in the normal state (**h**) or in the state of the PI3K/AKT pathway suppressed by LY294002 treatment (**i**). Data are shown as the mean ± s.d. of n = 3 cultures per condition. **a**, **e**, **f**, **h**, **i**, Significance (*p <0.05, **p <0.01, ***p <0.001, or ****p <0.0001) was calculated using Student’s *t*-test (**h**) or Dunnett’s test (**a**, **e**, **f**, **i**) for multiple comparisons.

*Mkl2*-KO mice also showed decreased skeletal muscle weight in the gastrocnemius, tibialis anterior, soleus, plantaris, and extensor digitorum longus muscles (21-22 weeks old, Fig. 1d, f, Extended Data Fig. 1e-l). Thus, the reduction in body weight of *Mkl2*-KO mice was mainly due to a reduction in the WAT and skeletal muscle mass (Fig. 1c-f, Extended Data Fig. 1a-l).

### Mkl1/2 repress Foxo1/3/4 transcriptional activity, independent of the PI3K/AKT signaling pathway

The loss of both WAT and skeletal muscle seen in *Mkl2*-KO mice was reminiscent of the phenotype caused by FoxO overactivation. Among FoxO transcription factors, *Foxo1* is a dominant regulator of adipocyte differentiation^12^. Therefore, we first investigated whether or not *Mkl2* deletion was associated with Foxo1 overactivation in inguinal WAT tissue.

We extracted RNA from inguinal WAT of *Mkl2*-KO mice and their littermate wild-type mice and analyzed their gene expression profiles. The resulting list of differentially expressed genes was analyzed using Ingenuity Pathway Analysis software to predict changes in the activation or inactivation status of transcription factors in *Mkl2*-KO mice (Fig. 1g, Supplementary Table 1). The transcriptional activity of Srf, a transcription factor that interacts with Mkl2, was inhibited in *Mkl2*-KO mice (Fig. 1g). Higher on the list than Srf was Foxo1, which was predicted to be in an activated state in *Mkl2*-KO mice. Although the expression of several Foxo1 target genes tended to increase, there was no marked change in *Foxo1* mRNA levels in the WAT of *Mkl2*-KO mice (Extended Data Fig. 1m), indicating that the transcriptional activity of Foxo1 was upregulated in the WAT of *Mkl2*-KO mice.

In addition to *Foxo1*, *Foxo3* and *Foxo4* are also involved in muscle atrophy^16^. Therefore, we further investigated whether or not Mkl2 could repress the transcriptional activity of Foxo1, Foxo3, and Foxo4 using a luciferase reporter with a FoxO-binding site. Mkl2 repressed the transcriptional activity of Foxo1, Foxo3, and Foxo4 (Fig. 1h). FoxO proteins are phosphorylated by AKT and then exported to the cytoplasm, where they are inactivated. Therefore, inhibition of the PI3K/AKT pathway inhibits FoxO phosphorylation and promotes FoxO activation by accumulating in the nucleus. Indeed, the transcriptional activities of Foxo1, Foxo3, and Foxo4 were further increased by treatment with LY294002, an inhibitor of PI3K (Fig. 1i). Interestingly, even when the PI3K/AKT pathway was inhibited, Mkl2 repressed the transcriptional activities of Foxo1, Foxo3, and Foxo4 (Fig. 1i). A similar inhibitory effect was observed for *Mkl1*, a paralog of *Mkl2* (Extended Data Fig. 2a, b). The repression of Foxo1/3/4 activity in the PI3K/AKT inhibited state by Mkl1/2 suggests that Mkl1 and Mkl2 are repressors of Foxo1/3/4, independent of the PI3K/AKT pathway.

### Foxo1 repression by Mkl2 relies on direct interaction, not post-translational modification

We next investigated whether or not the repression of Foxo1/3/4 by Mkl1/2 is dependent on protein-protein interactions. Co-immunoprecipitation experiments confirmed the binding of Foxo1/Mkl2 and Foxo1/Mkl1 (Fig. 2a and b). A split luciferase assay also confirmed the binding of all combinations of Foxo1/3/4 and Mkl1/2 (Fig. 2c, d, Extended Data Fig. 2c-f). We then performed split-luciferase experiments using a series of Foxo1 deletion constructs to determine which region of the Foxo1 protein binds to Mkl2. Foxo1, lacking the forkhead domain, showed significantly reduced binding to Mkl2 (Fig. 2e). The forkhead domain is required for Foxo1 to bind cis-regulatory elements in genomic DNA in the nucleus. Thus, Mkl2 may repress the transcriptional activity of Foxo1 by inhibiting its ability to bind DNA. When the nuclear translocation of Mkl1/2 was inhibited using CCG-1423^35^, the binding of Mkl1/2 to Foxo1 was significantly reduced, suggesting that Mkl1/2 binds to Foxo1 in the nucleus (Fig. 2f, Extended Data Fig. 2g). Split luciferase experiments using a series of *Mkl2* deletion constructs showed that the transactivation domain (TAD) of Mkl2 alone still binds to Foxo1 (Fig. 2g). Indeed, the TAD domain of Mkl2 alone was able to repress the transcriptional activity of Foxo1 (Fig. 2h). Furthermore, the TAD domain showed an inhibitory effect on the fusion protein of the forkhead domain of Foxo1 and VP16 (Fig. 2i), indicating that the TAD domain of Mkl2 binds to the DNA-binding domain of Foxo1 and inhibits its transcriptional activity of Foxo1. Thus, repression of FoxO transcriptional activity by Mkl1/2 is not achieved by post-translational modifications but by repressive interactions with the DNA-binding domain of FoxO.

**Fig. 2:**
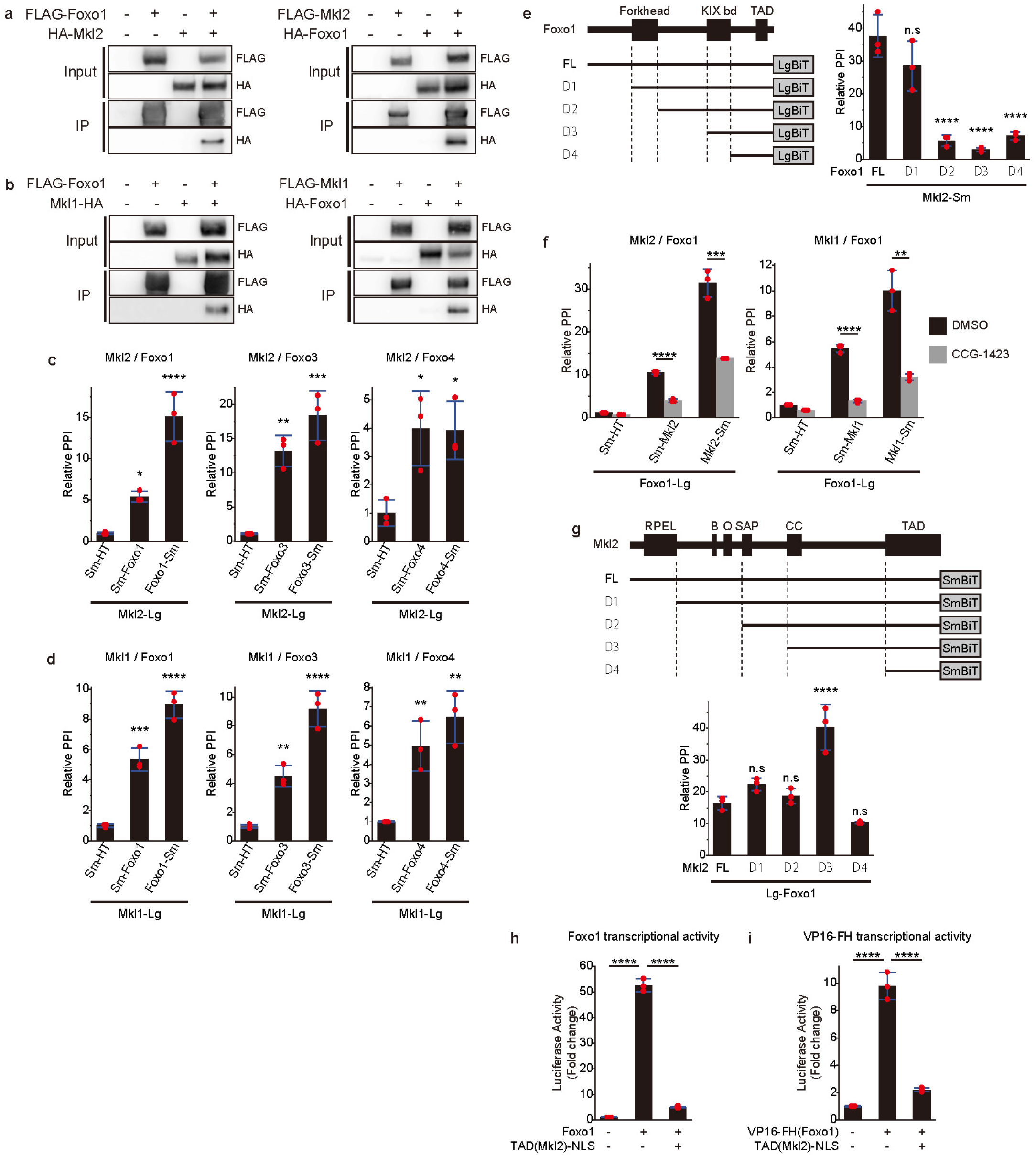
Mkl1 and Mkl2 bind directly to FoxO proteins. **a, b**, A co-immunoprecipitation assay of Foxo1 and either Mkl2 (**a**) or Mkl1 (**b**). FLAG-and HA-tagged proteins were expressed in 293FT cells. FLAG-tagged proteins were immunoprecipitated using FLAG antibody. Co-immunoprecipitation was confirmed by detection with HA antibody. **c, d**, Protein-protein interaction (PPI) of Foxo1, Foxo3, and Foxo4 with Mkl2 (**c**) or Mkl1 (**d**) was analyzed using split luciferase (NanoBiT). The relative PPI signal was calculated based on the signal between the LgBiT-tagged protein and SmBiT-Halotag (Lg: LgBiT tag, Sm: SmBiT tag, HT: Halotag). **e**, PPI of Mkl2 with truncated Foxo1 proteins using split luciferase (NanoBiT). **f**, Changes in the interaction of Foxo1 with Mkl2 or Mkl1 upon addition of CCG-1423, an inhibitor of the nuclear translocation of Mkl2 and Mkl1. **g**, PPI of Foxo1 with truncated Mkl2 proteins. **h, i**, Luciferase assay using a reporter with FoxO-responsive elements. Transcriptional activity of Foxo1 (**h**) or forkhead domain of Foxo1 and VP16 fusion protein (**i**) with or without nuclear-localized TAD domain of Mkl2. **c-i**, Data are shown as the mean ± s.d. of n = 3 cultures per condition. **c**-**i**, Significance (*p <0.05, **p <0.01, ***p <0.001, or ****p <0.0001) was calculated using Student’s *t*-test (**f**) or Dunnett’s test (**c**, **d**, **e**, **g**, **h**, **i**) for multiple comparisons. n.s: not significant.

### Mkl1/2 regulates Foxo1 activity in preadipocytes

The reduction in WAT observed in *Mkl2*-KO mice was investigated from the perspective of defects in preadipocyte differentiation due to Foxo1 overactivation^12^. Foxo1 represses both the initiation of *Pparγ* expression and its transcriptional activity^9–11^. Knockdown of *Mkl2* in 3T3-L1 preadipocytes using siRNA significantly impaired differentiation into mature adipocytes and suppressed the expression of *Pparγ* and white adipose differentiation markers (Fig. 3a-c). Furthermore, the transcriptional activity of Pparγ was repressed by Foxo1, but this repression was rescued by Mkl2 (Extended Data Fig. 3a). Mkl1 also showed a similar effect on Pparγ activity and adipocyte differentiation (Extended Data Fig. 3b-f). Adenoviral overexpression of *Foxo1* in 3T3-L1 preadipocytes suppressed adipocyte differentiation. However, co-expression of *Mkl2* or *Mkl1* partially rescued the differentiation defects caused by *Foxo1* overexpression (Fig. 3d). Similar to treatment with LY294002, a PI3K inhibitor, adipocyte differentiation was significantly suppressed in the presence of CCG-1423^35^, which inhibits Mkl1/2 nuclear translocation (Fig. 3e). *Pparγ* expression was also suppressed in the presence of CCG-1423 (Fig. 3f), indicating that Mkl1/2 regulates preadipocyte differentiation by repressing Foxo1 activity in the nucleus.

**Fig. 3:**
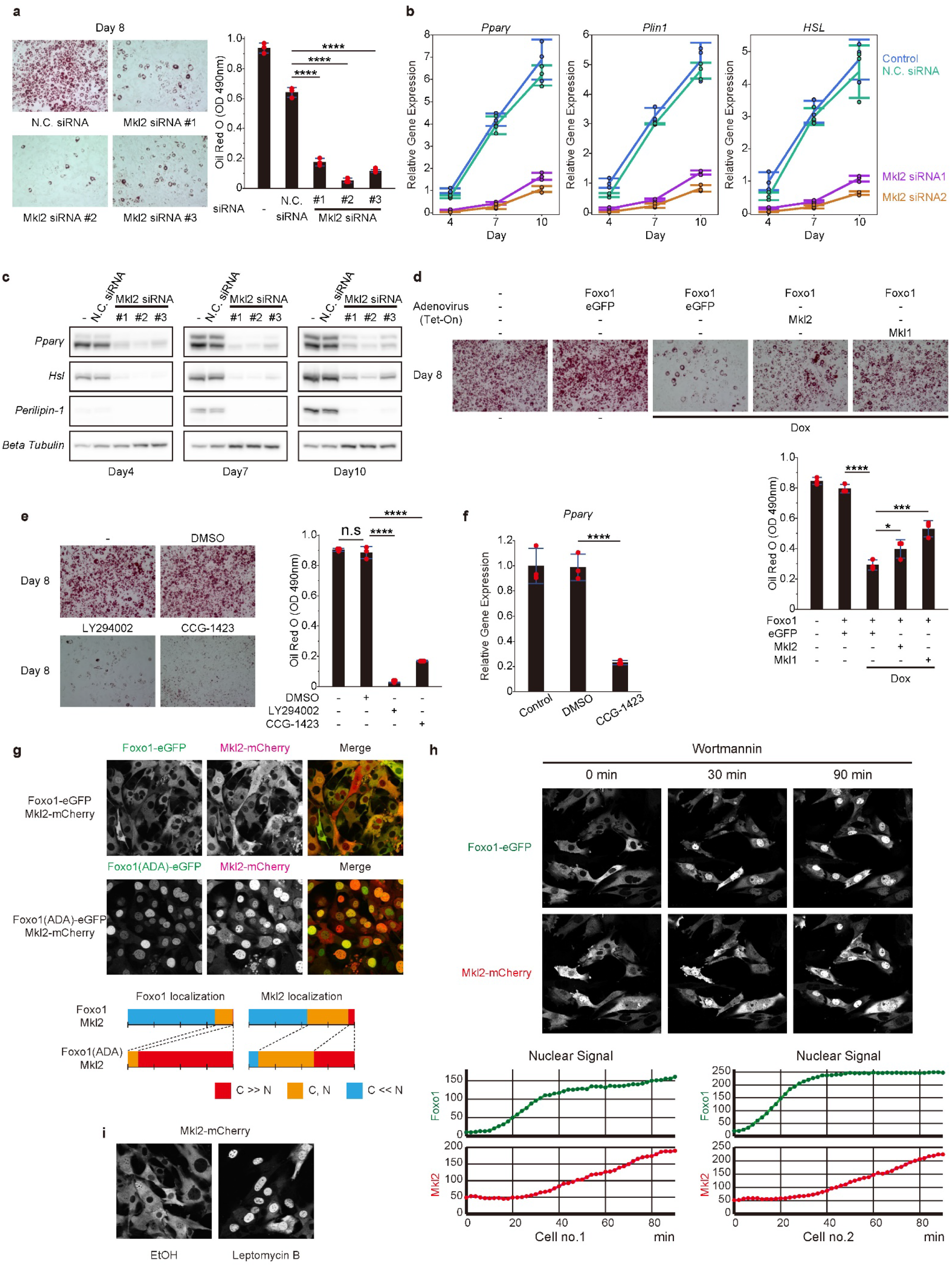
Inhibition of *Mkl2* significantly suppressed white adipocyte differentiation. **a**, Oil Red O staining of *Mkl2* knockdown 3T3-L1 cells at day 8 of differentiation. After staining, the dye was extracted with isopropanol and quantified by measuring absorbance at 490 nm (right). **b, c**, Expression of *Pparγ*, *Plin1* and *Hsl* mRNA (**b**) and protein (**c**) at days 4, 7, and 10 after induction of differentiation of 3T3-L1 cells with or without *Mkl2* knockdown. mRNA expression is shown as the relative change from the average expression of the control on day 4. **d**, Oil Red O staining of 3T3-L1 cells on day 8 of differentiation overexpressing *Foxo1*, *eGFP*, *Mkl1*, or *Mkl2*. After staining, the dye was extracted with isopropanol and quantified by measuring absorbance at 490 nm (bottom). **e**, Oil Red O staining of 3T3-L1 cells on day 8 of differentiation treated with LY294002 or CCG-1423. After staining, the dye was extracted with isopropanol and quantified by measuring absorbance at 490 nm (right). **f**, *Pparγ* mRNA expression in CCG-1423 treated 3T3-L1 cells on day 4 of differentiation. **g**, Subcellular localization of Mkl2 with Foxo1 or Foxo1(ADA) expression. The percentage of cells showing nuclear and cytoplasmic localization of Foxo1 and Mkl2 are shown below (C: cytoplasmic localization; N: nuclear localization). **h**, Localization changes of Foxo1 and Mkl2 after addition of wortmannin, a PI3K inhibitor. Changes in the nuclear signal over time are shown below for the two cells with representative behavior. **i**, Changes in Mkl2 localization after treatment with or without leptomycin B. **a, b, d-f**, Data are shown as the mean ± s.d. of n = 3 cultures per condition. Significance (*p <0.05, *p <0.001, or ****p <0.0001) was calculated using the Student’s *t*-test (**f**) or Dunnett’s test (**a**, **d**, **e**) for multiple comparisons.

Next, we investigated how Mkl1/2 responds to Foxo1 activation by analyzing its subcellular localization. Adenoviral expression of *Foxo1* and *Mkl2* fused to fluorescent proteins in 3T3-L1 preadipocytes showed that both Foxo1 and Mkl2 were mainly localized in the cytoplasm (Fig. 3g). However, constitutively active mutants of Foxo1(ADA) alter their localization to the nucleus. Surprisingly, although the Mkl2 protein remained wild type, its localization changed to the nucleus. Thus, in response to the localization of Foxo1, the subcellular localization of Mkl2 was determined (Fig. 3g). Similar localization changes in response to Foxo1 activity were also observed for Mkl1 (Extended Data Fig. 3g).

Time-lapse imaging was performed to visualize localization changes upon Foxo1 activation over time. When the PI3K inhibitor wortmannin was added, Foxo1 was rapidly translocated to the nucleus, followed by increased nuclear localization of Mkl2 (Fig. 3h, Supplementary Movie 1). Similar changes in Mkl2 and Mkl1 localization were observed with LY294002, a PI3K inhibitor (Extended Data Fig. 4a, b). In addition, both Mkl1 and Mkl2 accumulated in the nucleus when leptomycin B inhibited nuclear export (Fig. 3i, Extended Data Fig. 4c). These results suggest that Mkl1 and Mkl2 are constantly shuttling between the cytoplasm and nucleus, and once Foxo1 is activated, they remain in the nucleus and modulate Foxo1 activity.

### Atrogenes and *Mkl1* expression are always inversely correlated in various muscle atrophies

In skeletal muscle, various factors promote the nuclear translocation of Foxo1, Foxo3, and Foxo4 and strongly induce the expression of atrogenes, such as *atrogin-1* and *MuRF1*, leading to muscle protein degradation and even muscle atrophy^13–16^. Therefore, we investigated whether or not *Mkl1/2* is involved in skeletal muscle homeostasis by regulating the activities of Foxo1, Foxo3, and Foxo4.

First, we analyzed the subcellular localization of Mkl1/2 in C2C12 myotubes using adenoviral expression of *Mkl1* or *Mkl2* fused to eGFP. Mkl1 was exclusively localized to the nucleus, whereas Mkl2 was localized to the cytoplasm (Fig. 4a). This suggests that Foxo1/3/4 activity in the skeletal muscle may be under constant repression by Mkl1. We further investigated the changes in *Mkl1/2* during the onset of muscle atrophy. We examined various muscle atrophies with different causes, such as aging, sciatic denervation, fasting, steroid administration, and cancer cachexia. During aging, the expression of FoxO target genes, such as *atrogin-1* and *MuRF1*, significantly increased, while the expression of *Mkl1* decreased (Fig. 4b). The expression of *Mkl2* did not change with age. The expression of *Foxo1* and *Foxo3* increased, but the expression of *Foxo4* remained unchanged.

**Fig. 4:**
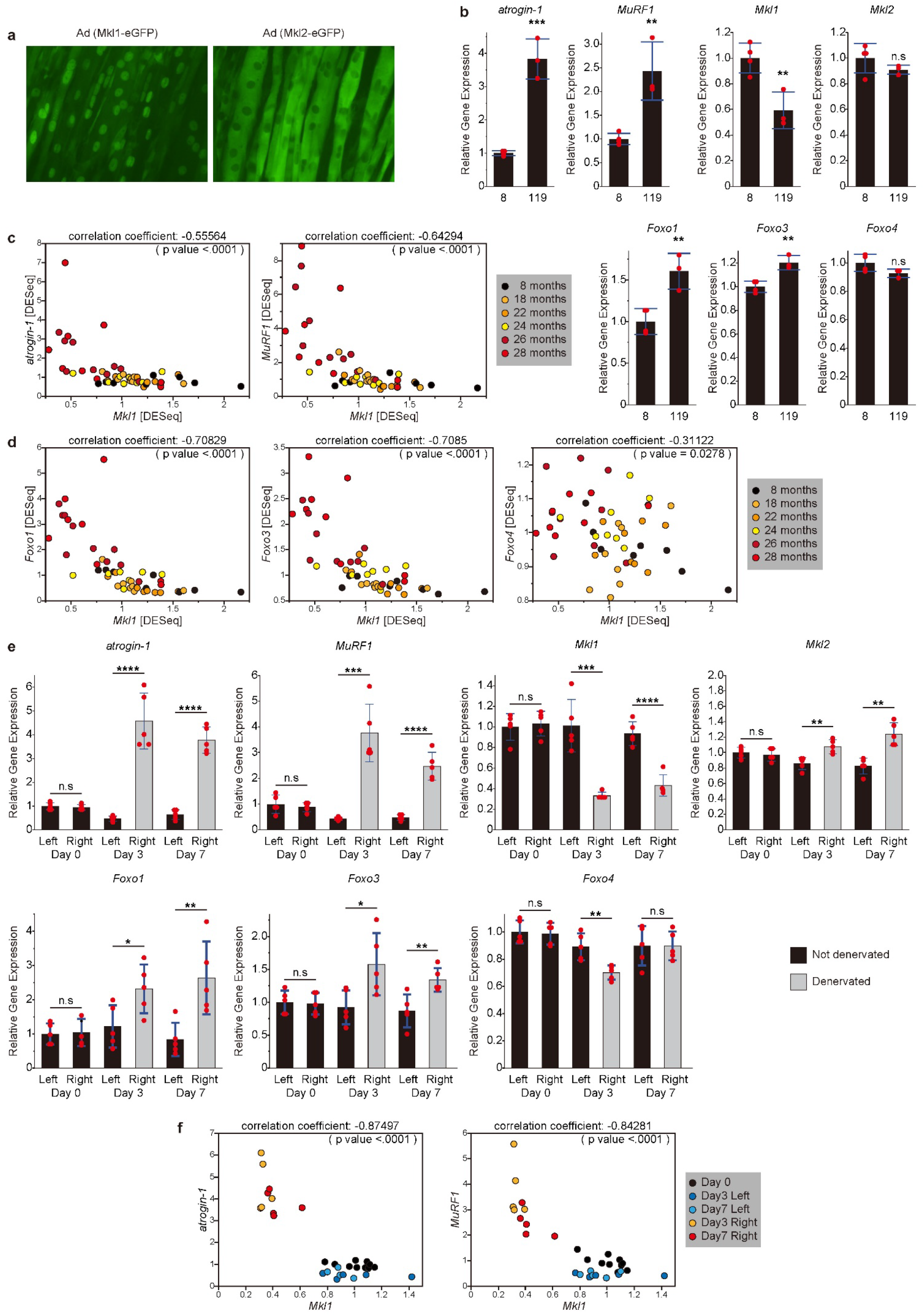
Expression changes of *atrogin-1*, *MuRF1*, *Foxo1/3/4*, and *Mkl1/2* in muscle atrophy induced by aging or sciatic nerve denervation. **a**, Mkl1-eGFP and Mkl2-eGFP were expressed in myotubes derived from C2C12 cells, and the subcellular localization of each protein was analyzed. To stabilize the protein levels, myotubes were treated with MG-132. **b**, Altered expression of *atrogin-1*, *MuRF1*, *Mkl1*, *Mkl2*, *Foxo1*, *Foxo3*, and *Foxo4* in the gastrocnemius muscle of young (8 weeks old, n=4) and old (119 weeks old, n=3) male C57BL/6J mice. **c, d**, Inverse correlation between the expression of *Mkl1* and atrogenes (**c**) or *Foxo1/3/4* (**d**) in gastrocnemius muscle of mice of different ages. Each point represents the DESeq-normalized gene expression in the gastrocnemius muscle of one mouse. These data were obtained from a re-analysis of the GEO dataset (GEO accession number: GSE145480). **e**, Altered expression of *atrogin-1*, *MuRF1*, *Mkl1*, *Mkl2*, *Foxo1*, *Foxo3*, and *Foxo4* in the left or right tibialis anterior muscle of mice with sciatic nerve denervation of the right hindlimb. Day = number of days after sciatic nerve denervation (n=5). **f**, Inverse correlation of *Mkl1* and atrogene expression in the tibialis anterior muscle of mice with right hindlimb sciatic nerve denervation. Each point represents the relative gene expression in either the left or right mouse tibialis anterior muscle. Relative gene expression was calculated based on the average expression in the left leg before sciatic nerve denervation (Day 0). **b, e,** Data are shown as the mean ± SD. Significance (*p <0.05, **p <0.01, ***p <0.001, or ****p <0.0001) was calculated using Student’s *t-*test. n.s: not significant.

We also re-analyzed comprehensive RNA-seq data from the mouse gastrocnemius muscle of 8-, 18-, 22-, 24-, 26-, and 28-month old mice^36^. Mice are thought to spontaneously develop sarcopenia around 26 months old. Indeed, the expression of *atrogin-1*, *MuRF1*, *Foxo1,* and *Foxo3* was significantly upregulated in mice after this age, while *Mkl1* expression was significantly downregulated (Extended Data Fig. 5a). These data also revealed a strong inverse correlation between atrogene expression and *Mkl1* (Fig. 4c). More surprisingly, between *Mkl1* and *Foxo1* or *Foxo3*, an even stronger inverse correlation was observed (Fig. 4d). As *Foxo4* expression increased only slightly with age, a weak inverse correlation with *Mkl1* was observed (Fig. 4b, d, Extended Data Fig. 5a).

Similarly, inverse correlations between atrogene and *Mkl1* expression have been observed in other muscle atrophy models. In sciatic nerve denervation-induced muscle atrophy, a significant increase in the expression of atrogenes was observed in the tibialis anterior and gastrocnemius muscles on the side where the sciatic nerve was denervated three and seven days after treatment. In contrast, *Mkl1* expression was significantly reduced in these muscles (Fig. 4e, Extended Data Fig. 5b). Strong inverse correlations were observed between atrogene and *Mkl1* expression in this model (Fig. 4f, Extended Data Fig. 5c). In addition, increased expression of *Foxo1* and *Foxo3*, but not *Foxo4*, was observed (Fig. 4e).

Similar inverse correlations between Mkl1 and atrogene, Foxo1, and Foxo3 expression were also observed in muscle atrophy induced by fasting, dexamethasone administration, and cancer cachexia. When mice were fasted for 24 h, increased expression of atrogenes, *Foxo1,* and *Foxo3* was observed in the gastrocnemius muscle, while *Mkl1* expression was significantly decreased (Fig. 5a, b). Similar changes in gene expression were also observed in mouse muscle as early as 16 h after intraperitoneal administration of dexamethasone (Fig. 5c). We also examined gene expression changes in skeletal muscle associated with cancer cachexia and found similar inverse correlations between *Mkl1* and atrogenes, *Foxo1,* and *Foxo3* (Fig. 5d). This analysis was performed by reanalyzing the data stored in the Gene Expression Omnibus (GEO) database^37^. Thus, in all of the muscle atrophy models examined, the expression of *Mkl1* was always repressed after the onset of muscle atrophy.

**Fig. 5:**
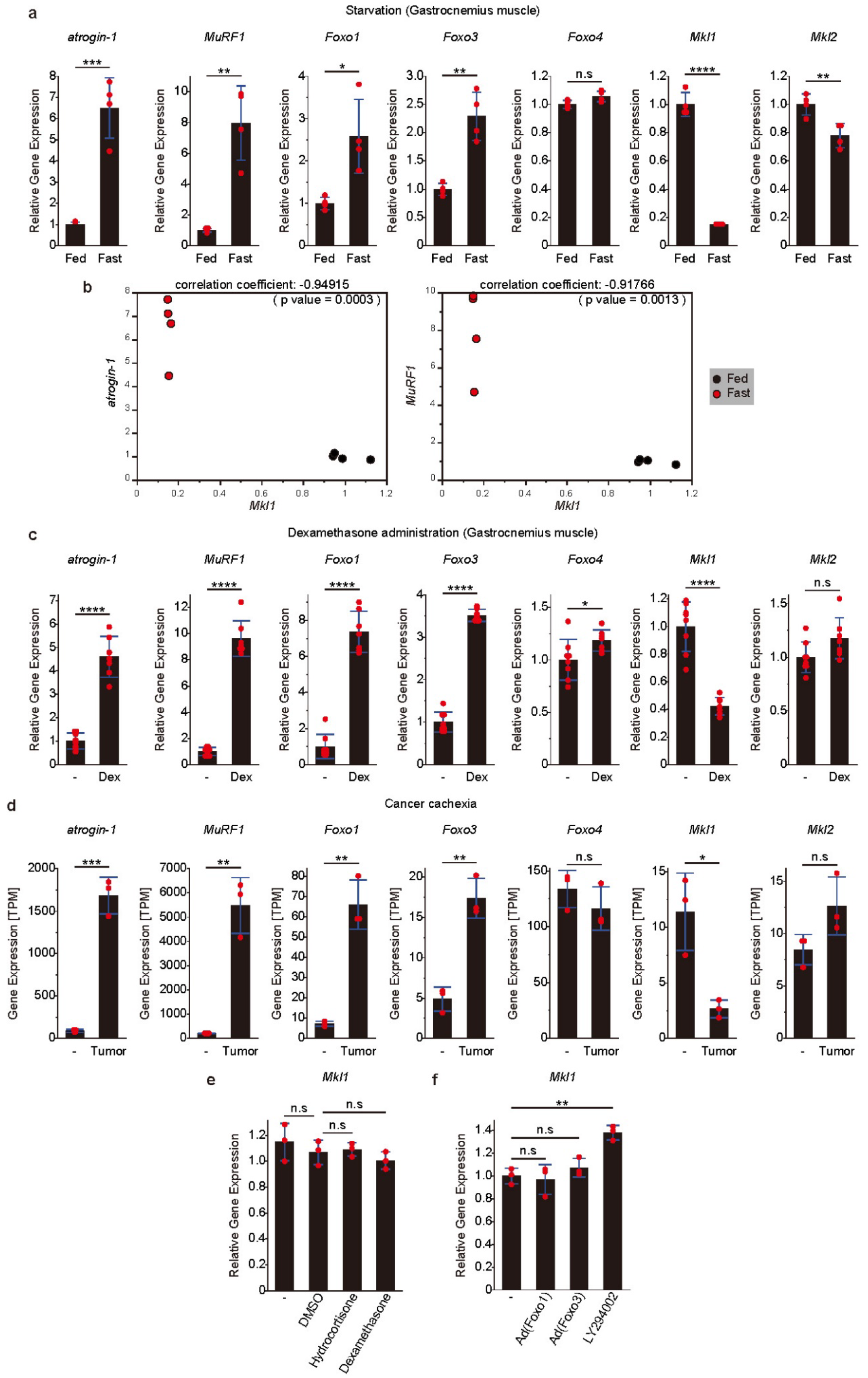
Expression changes of *atrogin-1*, *MuRF1*, *Foxo1/3/4*, and *Mkl1/2* in muscle atrophy induced by fasting, dexamethasone administration, or cancer cachexia. **a**, Altered expression of the *atrogin1*, *MuRF1*, *Foxo1*, *Foxo3*, *Foxo4*, *Mkl1*, and *Mkl2* in the gastrocnemius muscle of 9-week-old C57BL/6J male mice (n=4) fasted for 24 h. **b**, Inverse correlation between *Mkl1* and atrogene expression in the gastrocnemius muscle of mice fasted for 24 h. Each point represents relative gene expression in the gastrocnemius muscle of each mouse. Relative gene expression was calculated based on the average expression in fed mice. **c**, Altered expression of *atrogin1*, *MuRF1*, *Foxo1*, *Foxo3*, *Foxo4*, *Mkl1*, and *Mkl2* in the gastrocnemius muscle of C57BL/6J mice (n=7-8) 16 h after intraperitoneal administration of dexamethasone sodium phosphate. **d**, Altered expression of *atrogin1*, *MuRF1*, *Foxo1*, *Foxo3*, *Foxo4*, *Mkl1*, and *Mkl2* in mice (n=3) with cancer cachexia. These data were obtained from a re-analysis of the GEO dataset (GEO accession number: GSE65936). **e** Endogenous *Mkl1* expression in C2C12-derived myotubes treated with hydrocortisone or dexamethasone (n=3). **f**, Endogenous *Mkl1* expression in C2C12-derived myotubes when *Foxo1* or *Foxo3* was expressed by adenovirus or when Foxo1/3/4 was activated by the PI3K inhibitor LY294002 (n=3). **a**, **c**-**f**, Data are shown as the mean ± s.d. Significance (*p <0.05, **p <0.01, ***p <0.001, or ****p <0.0001) was calculated using Student’s *t*-test (**a**, **c**, **d**) or Dunnett’s test (**e**, **f**) for multiple comparisons.

To determine whether or not the decreased expression of *Mkl1* was a consequence of the activation of the muscle atrophy signaling pathway, we examined endogenous *Mkl1* expression during muscle catabolism (Fig. 5e, f). In C2C12 myotubes, addition of hydrocortisone, dexamethasone, or adenoviral overexpression of *Foxo1* and *Foxo3* did not reduce endogenous *Mkl1* expression. PI3K inhibition also failed to reduce *Mkl1* expression. These data suggest that the suppression of endogenous *Mkl1* expression may occur independently of activation of the GR/FoxO pathway or inhibition of the PI3K/AKT pathway.

### Mkl1 and Mkl2 repress the induction of atrogene expression in the skeletal muscle

We investigated whether or not *Mkl1* and *Mkl2* could inhibit FoxO-induced muscle atrophy in cultured myotubes. Since little upregulation of *Foxo4* was observed in the examined muscle atrophy models, we used adenoviral expression of *Foxo1* or *Foxo3* in myotubes to induce muscle atrophy. Overexpression of *Foxo1* or *Foxo3* in myotubes increased the expression of atrogenes, such as *atrogin-1* and *MuRF1* (Fig. 6a-d). This induction was suppressed by *Mkl1* or *Mkl2*, even in the presence of the PI3K inhibitor LY294002. Notably, *Mkl1* had a stronger inhibitory effect on atrogene expression than *Mkl2*, which may reflect differences in the subcellular localization of Mkl1 and Mkl2 in skeletal muscle cells (Fig. 4a). As mentioned above, Mkl2 binds to the forkhead domain of Foxo1 (Fig. 2e). Therefore, we performed chromatin immunoprecipitation of Foxo3 in myotubes to determine whether or not FoxO binding to the cis-regulatory region of downstream atrogenes was altered in the presence of Mkl1 or Mkl2. From the ChIP-Atlas database^38^, regions with high scores in the previous Foxo3 ChIP-seq data were selected as Foxo3 binding sites. Foxo3 binding was observed at these sites near *atrogin-1* and *MuRF1*, but co-expression of *Mkl1* or *Mkl2* reduced Foxo3 binding at these sites (Extended Data Fig. 6). This suggests that Mkl1 and Mkl2 bind to Foxo3 to inhibit DNA binding in the skeletal muscle.

**Fig. 6:**
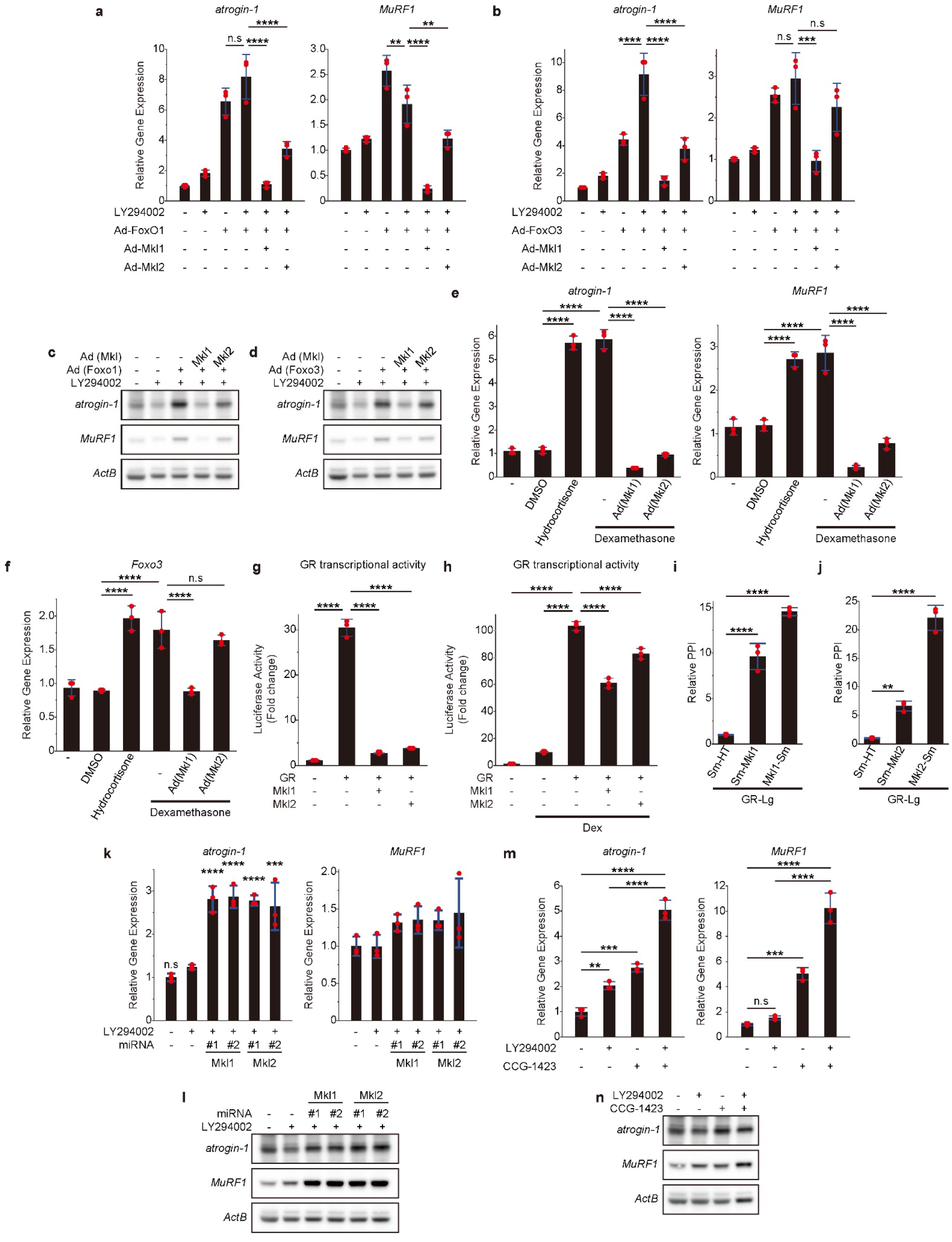
Mkl1 and Mkl2 suppressed the induction of atrogene expression in skeletal muscle. **a, b**, Inhibitory effects of *Mkl1* or *Mkl2* on mRNA expression of atrogenes induced by *Foxo1* (**a**) or *Foxo3* (**b**) in C2C12-derived myotubes. Adenoviruses were used to express *Foxo1*, *Foxo3*, *Mkl1*, or *Mkl2*. Ad: Adenovirus vector. **c, d**, Inhibitory effects of *Mkl1* or *Mkl2* on protein expression of atrogenes induced by *Foxo1* (**c**) or *Foxo3* (**d**) in C2C12-derived myotubes. Adenoviruses were used to express *Foxo1*, *Foxo3*, *Mkl1*, or *Mkl2*. Ad: Adenovirus vector. **e, f**, Inhibitory effect of *Mkl1* or *Mkl2* on atrogene (**e**) and *Foxo3* (**f**) mRNA expression induced by glucocorticoid analogs in C2C12-derived myotubes. An adenovirus was used to express *Mkl1* or *Mkl2*. Ad: Adenovirus vector. **g, h**, Inhibitory effect of *Mkl1* and *Mkl2* on the transcriptional activity of the glucocorticoid receptor (GR) using luciferase reporters with GR-responsive elements in the absence (**g**) or presence (**h**) of dexamethasone. **i, j** Protein-protein interaction (PPI) between GR and Mkl1 (**i**) or Mkl2 (**j**) detected using split luciferase (NanoBiT). The relative PPI signal was calculated based on the signal between the LgBiT-tagged GR and SmBiT-Halotag (Lg: LgBiT tag, Sm: SmBiT tag, HT: Halotag). **k, l,** Changes in mRNA (**k**) or protein (**l**) expression of atrogenes when *Mkl1* or *Mkl2* was suppressed in C2C12-derived myotubes. Knockdown was achieved by infection with adenoviruses expressing artificial microRNAs (miRNAs) targeting *Mkl1* or *Mkl2*. **m, n,** Altered mRNA (**m**) or protein (**n**) expression of atrogenes in C2C12-derived myotubes in the presence of the PI3K inhibitor LY294002 and/or the Mkl1/2 nuclear translocation inhibitor CCG-1423. **a, b, e-k, m**, Data are shown as the mean ± s.d. of n = 3 cultures per condition. Significance (**p <0.01, ***p <0.001, or ****p <0.0001) was calculated using Dunnett’s test (**a**, **b**, **g**-**k**) or Tukey’s test (**e**, **f**, **m**) for multiple comparisons. n.s: not significant.

Similarly, the expression of atrogenes induced by glucocorticoids was suppressed by *Mkl1* or *Mkl2* (Fig. 6e). Surprisingly, *Mkl1* also blocked the induction of *Foxo3* expression by GR activation (Fig. 6f). We found that Mkl1 and Mkl2 directly bound to GR and repressed its transcriptional activity (Fig. 6g-j). Thus, Mkl1 and Mkl2 repress both FoxO and GR activity in the GR/FoxO axis.

In contrast to the overexpression of *Mkl1* or *Mkl2*, knockdown of *Mkl1* or *Mkl2* resulted in increased expression of *atrogin-1* and *MuRF1* (Fig. 6k, l). Although *MuRF1* mRNA expression changes following *Mkl1* or *Mkl2* knockdown were relatively small, significant upregulation was observed at the protein level. Similarly, inhibition of nuclear localization of Mkl1 and Mkl2 by CCG-1423 alone resulted in increased expression of *atrogin-1* and *MuRF1*, and dual inhibition of Mkl1/2 nuclear localization and the PI3K/AKT pathway with CCG-1423 and LY294002 synergistically and strongly increased the expression of atrogenes (Fig. 6m, n). This result indicated that the suppression of Foxo1/3/4 by Mkl1/2 is independent of the PI3K/AKT pathway.

### Mkl1/2 supplementation inhibits skeletal muscle atrophy

To test the protective effects of *Mkl1* and *Mkl2* against muscle atrophy, we examined the size of the C2C12 myotubes. Overexpression of *Foxo1* or *Foxo3* caused severe atrophy of myotubes. However, when *Mkl1* or *Mkl2* were co-expressed with *Foxo1* or *Foxo3*, no such atrophy was observed (Fig. 7a). Based on the fact that *Mkl1* and *Mkl2* have repressive effects on FoxO transcriptional activity and that *Mkl1* is drastically reduced at the onset of muscle atrophy, we hypothesized that if *Mkl1* expression could be compensated for during muscle atrophy, it might be possible to confer resistance to skeletal muscle atrophy.

**Fig. 7:**
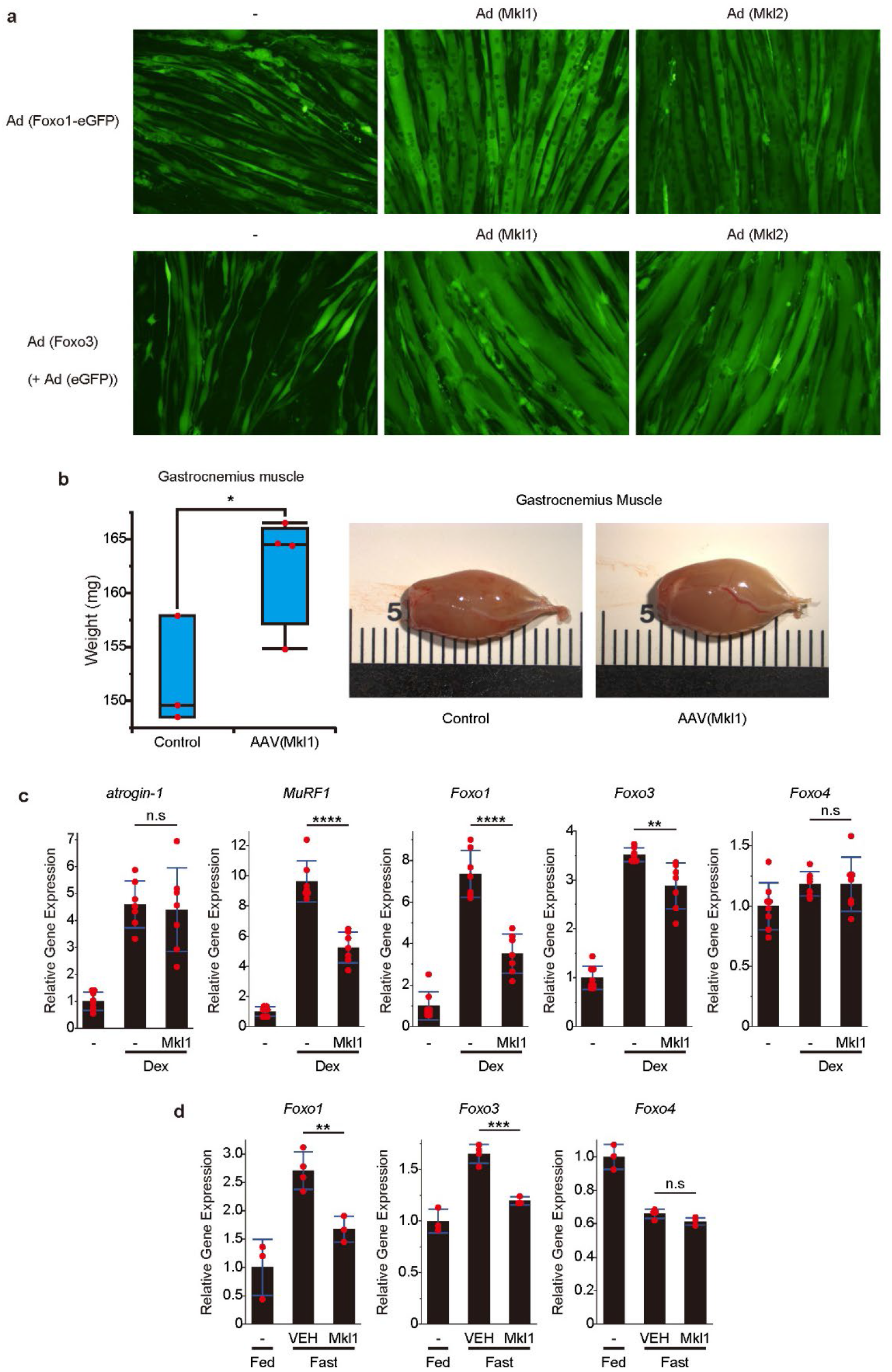
Mkl1 and Mkl2 inhibited skeletal muscle atrophy. **a**, Suppression of muscle atrophy by expression of *Mkl1* or *Mkl2* in C2C12-derived myotubes in which muscle atrophy was induced by the expression of *Foxo1* or *Foxo3*. Adenoviruses were used to express *Foxo1*, *Foxo3*, *Mkl1*, or *Mkl2* in myotubes. Ad: Adenovirus vector. **b**, Gastrocnemius muscle of mice 12 weeks after administration of 1×10^11^ vg of MyoAAV.2A (Mhck7-*Mkl1*) via tail vein. (Control: n=3, AAV: n=4) **c**, A comparison of gene expression of *atrogin-1*, *MuRF1*, *Foxo1*, *Foxo3*, and *Foxo4* in gastrocnemius muscle 16 h after administration of dexamethasone sodium phosphate (Dex). Dex was administered intraperitoneally to mice 12 weeks after MyoAAV.2A (Mhck7-*Mkl1*) administration. Data are shown as the mean ± s.d. (n= 7-8). **d**, A comparison of *Foxo1*, *Foxo3*, and *Foxo4* expression in the gastrocnemius muscle of mice expressing *Mkl1* in muscle by MyoAAV.2A and fasted for 24 h. Data are shown as the mean ± s.d. (n= 7-8). **b**-**d**, Significance (*p <0.05, **p <0.01, ***p <0.001, or ****p <0.0001) was calculated using Student’s *t*-test. n.s: not significant.

A modified adeno-associated virus (MyoAAV.2A) with strong muscle tropism has been reported^39^. Using MyoAAV.2A, we attempted to express *Mkl1* in the muscles throughout the body. First, MyoAAV.2A encoding *AkaLuc*-*eGFP* was systemically administered to mice, and its expression was confirmed. AkaLuc-eGFP was successfully induced in muscles, such as the gastrocnemius and tibialis anterior, as well as in the heart (Extended Data Fig. 7). The expression reached a plateau approximately two weeks after administration. We then administered MyoAAV.2A, which encodes *Mkl1*, to mice. Significant cardiac hypertrophy was observed following administration of AAV encoding *Mkl1* at doses greater than 2 × 10^11^ vg (Extended Data Fig. 8). Therefore, we had to administer lower doses of AAV to mice to assess skeletal muscle changes. After 10 weeks of AAV treatment, the muscles of mice treated with MyoAAV.2A, encoding *Mkl1*, were heavier than those of the untreated group (Fig. 7b).

We then treated the mice with dexamethasone and examined the changes in the expression of atrogenes and Foxo1/3/4. Dexamethasone-induced upregulation of *atrogin-1* could not be suppressed by treatment with MyoAAV.2A, which encodes *Mkl1*, but *MuRF1* and other atrogenes were suppressed (Fig. 7c). In addition, the dexamethasone-induced upregulation of *Foxo1* and *Foxo3* was blocked by *Mkl1* (Fig. 7c). We then examined whether or not fasting-induced expression of atrogenes was altered by *Mkl1* supplementation. In *Mkl1*-supplemented gastrocnemius muscle, the fasting-induced increase in *Foxo1* and *Foxo3* expression was significantly suppressed (Fig. 7d). Although the induction of *atrogin-1* and *MuRF1* expression was barely suppressed, the increased expression of several other atorogenes was significantly suppressed (Extended Data Fig. 9). Thus, *Mkl1* supplementation of skeletal muscle showed a protective effect against the induction of skeletal muscle atrophy.

### Exercise activates *Mkl1/2* by promoting nuclear accumulation of Mkl2 and increasing *Mkl1/2* expression in skeletal muscle

As mentioned above, *Mkl1* expression was invariably reduced at the onset of muscle atrophy (Fig. 4b, e; Fig. 5a, c, d; Extended Data Fig. 5a, b), reducing its inhibitory effect on GR and Foxo1/3/4. Finally, we investigated whether *Mkl1/2* is activated during the anabolic process or during the inhibition of catabolism in skeletal muscle. Exercise induces muscle hypertrophy and inhibits muscle atrophy. Therefore, we investigated whether or not skeletal muscle contraction and exercise affected *Mkl1/2*.

Mkl2 was localized in the cytoplasm of myotubes (Fig. 4a). Interestingly, the nuclear translocation of Mkl2 was observed in response to contraction in myotubes (Fig. 8a, Supplementary Movie 2). This suggests that myotube contraction increases the inhibitory effect of *Mkl2* on muscle atrophy in the skeletal muscle cells. Similarly, when the gastrocnemius muscle was excised immediately after treadmill exercise and the cellular proteins were fractionated, the amount of nuclear Mkl2 protein was significantly increased (Fig. 8b). In addition, we found that four weeks of treadmill exercise in mice increased the skeletal muscle expression of *Mkl1* and *Mkl2* (Fig. 8c), as in human skeletal muscle after long-term resistance training^40^. These results suggest that exercise activates *Mkl1/2* in skeletal muscle, both by increasing the nuclear accumulation of Mkl2 and by increasing the expression of *Mkl1* and *Mkl2*. Thus, some of the beneficial effects of exercise may involve *Mkl1* and *Mkl2*.

**Fig. 8:**
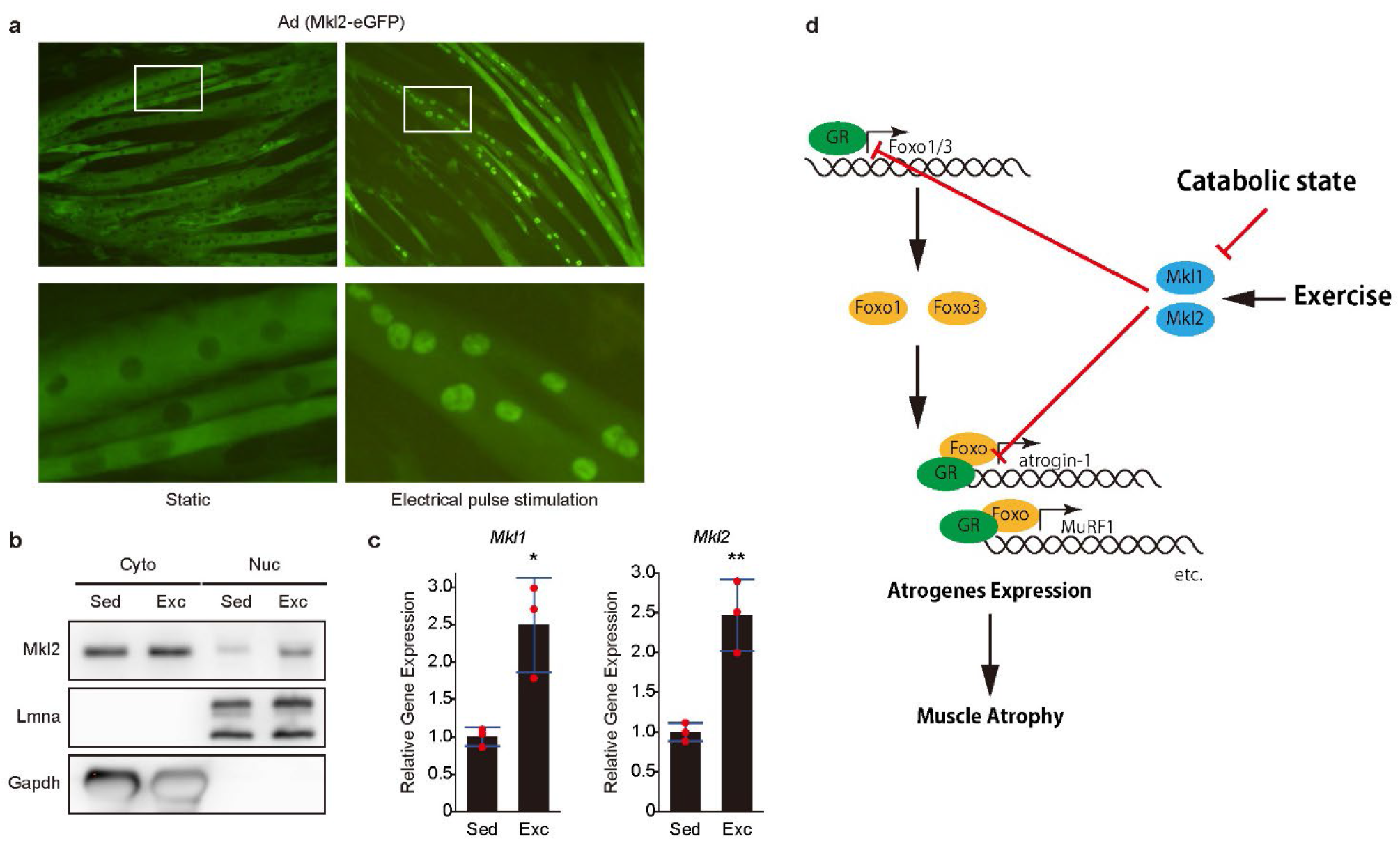
Exercise promotes the nuclear accumulation of Mkl2 and also increases the expression of both *Mkl1* and *Mkl2*. **a**, Changes in the subcellular localization of Mkl2 due to contraction of myotubes induced by electrical pulse stimulation (EPS). **b,** Subcellular localization of Mkl2 in the gastrocnemius muscle before and after treadmill exercise in C57BL/6J mice. **c**, Changes in *Mkl1* and *Mkl2* expression in the gastrocnemius muscle after 4 weeks of treadmill exercise in C57BL/6J mice (Sed: sedentary, n=3, Exc: exercise, n=3). Significance (*p <0.05, **p <0.01) was calculated using Student’s *t*-test. **d** A schematic diagram showing the effects of Mkl1 and Mkl2 on the GR/FoxO pathway, which regulates the progression of muscle atrophy. Exercise activates *Mkl1/2* through the nuclear translocation of Mkl2 and increased *Mkl1/2*. However, *Mkl1* expression decreases in response to the onset of muscle atrophy. GR: glucocorticoid receptor.

## DISCUSSION

In this study, we identified a novel mechanism of Foxo1/3/4 repression that is independent of the PI3K/AKT pathway. Both Mkl1 and Mkl2 are central to the repression mechanism. Mkl1/2 appears to repress the transcriptional activity of Foxo1/3/4 by inhibiting its ability to bind DNA. The activity of Foxo1/3/4 is thought to be primarily regulated by post-translational modifications, but the mode of repression by Mkl1/2 is based on direct protein-protein interactions. The regulation of Foxo1/3/4 activity by Mkl1/2 is important for the maintenance of at least two tissues: WAT and skeletal muscle. The plasticity of these two tissues is significantly affected by aging. Therefore, it is possible that the Mkl1/2 activity decreases with age. Indeed, we observed a significant decrease in *Mkl1* expression in the skeletal muscle of mice with age.

In preadipocytes, we demonstrated a novel mechanism by which Mkl1/2 rapidly responds to Foxo1 activation by shuttling between the cytoplasm and nucleus. When Foxo1 is activated and accumulates in the nucleus, Mkl1/2 remains in the nucleus, preventing Foxo1 overactivation, thereby ensuring *Pparγ* expression. In contrast, in skeletal muscle, myocyte contraction promotes nuclear translocation of Mkl2, and continuous exercise increases *Mkl1/2* expression, indicating that exercise activates *Mkl1/2*. Mkl1/2 inhibited the GR/FoxO pathway involved in skeletal muscle atrophy (Fig. 8d). Surprisingly, in all of the muscle atrophy models studied, there was a mechanism by which *Mkl1* expression was repressed with the onset of muscle atrophy. This suggests that downregulation of *Mkl1* is essential for the progression of muscle atrophy and that *Mkl1* is a potent inhibitor of muscle atrophy. Indeed, *Mkl1* showed a strong inhibitory effect on Foxo1/3/4 and GR.

*Mkl2* is clearly required for WAT plasticity, as a decrease in WAT was observed in *Mkl2*-KO mice. However, our results conflict with several reports on the effects of *Mkl1* on adipogenesis. Nobusue et al. reported that *Mkl1* negatively regulates white adipocyte differentiation by repressing the transcriptional activity of *Pparγ*^41^. McDonald et al. reported that the loss of *Mkl1* in mesenchymal stem cells promotes lineage commitment to brown adipocytes^42^. *Mkl1*-KO mice appeared to have reduced amounts of subcutaneous and visceral WAT. At the same time, the expression of thermogenic genes, such as *Ucp1*, was increased in subcutaneous WAT^42^. In contrast, our results show that Mkl1, like Mkl2, can also repress the transcriptional activity of Foxo1 and that knockdown of *Mkl1* in preadipocytes significantly impairs *Pparγ* expression and white adipocyte differentiation. Adipogenesis involves several steps, including multipotent mesenchymal precursors, preadipocyte commitment, growth arrest, clonal expansion, terminal differentiation, and maturation. During this process, Foxo1 switches between activated and inactivated states, and the onset of *Pparγ* expression corresponds to the second inactivation of Foxo1^43^. The role of *Mkl1* may change at each step, but at least with respect to the repression of *Pparγ* by *Foxo1*, *Mkl1* may be as necessary as *Mkl2* for white adipocyte differentiation. Incidentally, it has been reported that Foxo1 activation increases *Ucp1* expression in WAT^44^. Thus, the increased expression of *Ucp1* in *Mkl1*-KO mice may be partly due to the activation of Foxo1.

*Srf* is already known to be important for skeletal muscle development^45^. The present study showed that the cofactors *Mkl1* and *Mkl2* play a protective role against muscle protein degradation by suppressing Foxo1/3/4 and GR independently of *Srf*. Interestingly, *Srf* and its cofactors are responsible for skeletal muscle development and the inhibition of atrophy. In addition, *Mkl1* and *Mkl2* are activated by exercise. Exercise is effective in preventing sarcopenia, and it is now clear that *Mkl1* and *Mkl2* are involved in this molecular mechanism. In the future, the development of agents that can increase the expression of *Mkl1* or *Mkl2* or promote the nuclear localization of Mkl2 in skeletal muscle is expected to mimic exercise and inhibit muscle atrophy.

We used an adeno-associated virus (MyoAAV.2A) to induce *Mkl1* expression in the skeletal muscles of mice. There was an upper limit to the amount of virus that could be administered, as high levels of *Mkl1* expression in the myocardium caused cardiac hypertrophy. Nevertheless, despite the small amount of virus compared to other studies, *Mkl1* was able to suppress the induction of the expression of many atrogenes, *Foxo1* and *Foxo3* in skeletal muscles, resulting in muscle hypertrophy. A more potent inhibitory effect on muscle atrophy may be achieved using an adeno-associated virus with a more skeletal muscle-specific promoter.

Incidentally, the overactivation of FoxO proteins has been reported in dilated cardiomyopathy (DCM) caused by *Lmna* mutations^34^. The nuclear localization of Mkl1 is markedly attenuated in *Lmna*-deficient or N195K mutant cells^33^, suggesting that *Mkl1/2* dysfunction may be involved in the pathogenesis of DCM. *Foxo1* overactivation in the myocardium has also been implicated in the pathogenesis of diseases other than DCM. For example, doxorubicin, an anticancer drug, increases the transcriptional activity of Foxo1 in the myocardium, leading to myocardial atrophy^46^. Therefore, inhibition of Foxo1 by Mkl1 or Mkl2 may be a potential therapeutic strategy for anthracycline-induced cardiac injury.

## MATERIALS AND METHODS

### Mouse experiments

ES cells (MrtfbTM1a(KOMP)Mbp) were obtained from the Knockout Mouse Project (KOMP) repository and used to generate chimeric mice. MrtfbTM1a(KOMP)Mbp mice were sequentially crossed with CAG-FRT and CAG-Cre mice to generate heterozygous mice lacking exons 11 and 12 of *Mkl2*. Heterozygous mice were maintained in C57BL/6N and ICR mouse strains, and *Mkl2*-knockout mice lacking both alleles were generated by heterosis between the two strains. C57BL/6J mice (including aged mice) used in the other animal experiments were obtained from CLEA Japan Inc. Other aged mice were also provided by the Institute of Development, Aging, and Cancer (IDAC), Tohoku University, and the Kobe Biomedical Research Foundation under the National BioResource Project of the Ministry of Education, Culture, Sports, Science and Technology.

Long-term exercise training was performed as follows: 10-week-old C57BL/6J male mice were exercised on a treadmill for 60 min per day for 4 weeks. Forced running was started at a speed of 15 m/min, and the speed was increased by 1 m/min every 10 min until it reached 20 m/min. After four weeks of training, the mice were dissected and analyzed for gene expression in the gastrocnemius muscle. For subcellular fractionation of muscle proteins, gastrocnemius muscle from mice trained on a treadmill for 3 h was fractionated into the cytoplasm and nucleus using the Subcellular Protein Fractionation Kit for Tissues (Thermo Fisher Scientific).

In the denervation-induced muscle atrophy model, the sciatic nerve was denervated in the right hindlimb of 10-week-old C57BL/6J male mice, and the right and left gastrocnemius and tibialis anterior muscles were harvested 3 and 7 days after denervation. For fasting-induced muscle atrophy, 9-week-old C57BL/6J male mice were fasted for 24 h. After fasting, the mice were euthanized, and the gastrocnemius muscles were harvested.

Dexamethasone sodium phosphate was dissolved in phosphate-buffered saline (PBS) and administered intraperitoneally to mice. AAV was also dissolved in PBS and administered to the mice via the tail vein. Transgene expression after AAV administration was confirmed by measuring bioluminescence using the IVIS Lumina II (Caliper Life Sciences) or eGFP fluorescence.

The mouse experiments were approved by Institutional Laboratory Animal Care and Use Committee of Tohoku University and Safety Committee for Recombinant DNA Experiments of Tohoku University.

### Reagents and antibodies

The reagents used in this study were as follows: Dexamethasone, IBMX, and insulin solution were obtained from Sigma-Aldrich; hydrocortisone was obtained from PromoCell; CCG-1423 and leptomycin B were obtained from Cayman Chemical; MG-132, LY-294002, doxycycline hydrochloride n-hydrate, and dexamethasone sodium phosphate were purchased from Fujifilm Wako Chemicals.

The following antibodies were used in this study: anti-PPARγ (C26H12, #2435), anti-HSL (#4107), anti-perilipin-1 (D418, #3470), anti-lamin A/C (#2032), anti-HA tag (C29F4, #3724), and normal rabbit IgG (#2729) from Cell Signaling Technology; anti-atrogin-1 (F-9, sc-166806) and anti-MuRF1 (C-11, sc-398608) from Santa Cruz Biotechnology; anti-GAPDH (M171-7), anti-β-actin (PM053-7), anti-DDDDK-tag (PM020-7), and anti-HA-tag (561-7) from MBL; and anti-beta-tubulin (HRP-66240) from Proteintech.

The siRNAs used in this study (27mer DsiRNA) were the following predesigned products from Integrated DNA Technologies: *Mkl1* siRNA #1 (#106160699), *Mkl1* siRNA #2 (#106037572), *Mkl1* siRNA #3 (#106037569), *Mkl2* siRNA #1 (#105942460), *Mkl2* siRNA #2 (#105942457), *Mkl2* siRNA #3 (#105942454), and Negative Control DsiRNA (#51-01-14-04).

### Plasmids and cosmids

The *Foxo1/3/4* responsive luciferase reporter plasmid was constructed by inserting six FoxO REs (FoxO-responsive elements) into the pGL4.23 (Promega). The following sequence was used for the FoxO RE: 5*’*-TGTTTGGCCGCTCTGTTTACGGCAGCAAGAGGATCTGTTTTTGGCCGCTTGTTTACGGCAAGTAAACA-3 *’*. Similarly, a Pparγ-responsive luciferase reporter was created by incorporating three PPRE (Pparγ responsive elements) into the pGL4.23. The single PPRE sequence is 5*’*-AGGGGACCAGGACAAAGGTCACGTTCGGGA-3*’*. A glucocorticoid receptor-responsive luciferase reporter was created by incorporating four glucocorticoid receptor-responsive elements (GREs) into the pGL4.23. The single GRE sequence is 5*’*-GGTACATTTTGTTCT-3*’*. The second of the four GREs was placed in the antisense orientation.

Protein coding sequences (CDS) for *Foxo1*, *Foxo3*, *Foxo4*, and the glucocorticoid receptor (*Nr3c1*) were amplified by polymerase chain reaction (PCR) from mouse cDNA. The CDSs for *Mkl1* and *Mkl2* were synthesized as DNA fragments, and these codons were optimized (Integrated DNA Technologies). These CDSs were cloned into mammalian expression vectors (pcDNA3.1) or NanoBiT PPI starter system vectors (pBiT1.1-N, pBiT1.1-C, pBiT2.1-N, and pBiT2.1-C).

The Tet-On adenoviral cosmid vector was produced using the following procedure: the multicloning site of the pTetOne vector (TaKaRa) was modified to retain a SwaI recognition sequence. The entire transgene expression regulatory region of the pTetOne vector was inserted into the pAxcwit2 cosmid vector to create the pAxcwit2-TetOne vector. Finally, transgenes, such as *Foxo1/3/4* or *Mkl1/2* were inserted into pAxcwit2-TetOne using the SwaI site.

AAV transfer plasmids were constructed by removing all the sequences between the ITRs of the pAAV-CMV vector (TaKaRa) and inserting the Mhck7 promoter, transgene, and SV40 polyA sequences. DNA templates for the Mhck7 promoter, *AkaLuc* and *Mkl1* were prepared using gene synthesis (Integrated DNA Technologies). The Rep/Cap plasmid encoding the MyoAAV.2A capsid was generated by modifying pUCmini-iCAP-PHP. S by inverse PCR^47^. The hypervariable region VIII sequence SAQQAVRTSL was replaced with GPGRGDQTTL. pUCmini-iCAP-PHP.S was a gift from Viviana Gradinaru (Addgene plasmid #103006; http://n2t.net/addgene:103006; RRID:Addgene_103006). The helper plasmid used was pHelper (TaKaRa).

### Cell culture

3T3-L1 cells (ATCC CL-173) were maintained in DMEM (Nacalai Tesque) containing 10% calf serum (Gibco). For differentiation into adipocytes, 3T3-L1 cells were cultured to confluence, after which the medium was changed, and the cells were cultured for a further 48 h. To initiate differentiation, the medium was replaced with differentiation medium (90% DMEM, 10% FBS, 1 mM sodium pyruvate, 1 µM dexamethasone, 0.5 mM IBMX, and 1 µg/mL insulin). After 48 h, the differentiation medium was replaced with a maintenance medium (90% DMEM, 10% FBS, 1 mM sodium pyruvate, and 1 µg/mL insulin), which was then changed every 48 h.

C2C12 cells (ATCC CRL-1772) were seeded in dishes coated with Geltrex LDEV-Free Reduced Growth Factor Basement Membrane Matrix (Thermo Fisher Scientific) and maintained in DMEM (Nacalai Tesque) containing 10% FBS (Gibco). For differentiation into myotubes, the culture medium was replaced with DMEM containing 2% horse serum (Gibco) after the C2C12 cells reached confluence. Three days after the induction of differentiation, adenovirus was added to the medium to infect the myotubes, and the next day, doxycycline was added to induce transgene expression. 293FT cells were purchased from Thermo Fisher Scientific and maintained in DMEM (Nacalai Tesque) containing 10% FBS (Gibco).

### Electrical pulse stimulation

Electrical pulse stimulation was applied to the well-differentiated myotubes on day 5 of differentiation. Two graphite plates were inserted into the wells, and a constant-voltage bipolar square pulse was generated between the graphite plates at 1 Hz using a C-Pace EM (IonOptix) to contract the myotubes.

### Luciferase assays

Luciferase reporter and plasmids encoding each gene were transfected into 293FT cells (Thermo Fisher Scientific) using Polyethylenimine Max (molecular weight [MW]: 40,000; Polysciences). If necessary, LY294002 or dexamethasone was added 24 h after transfection. Forty-eight hours after transfection, cells were harvested and lysed. D-luciferin and ATP were added to the cell extract, and luminescence intensity was measured using a GloMax Discover Microplate Reader (Promega).

### Detection of protein-protein interactions

For co-immunoprecipitation assays, 293FT cells were transfected with plasmids encoding FLAG- and/or HA-tagged proteins using Polyethylenimine Max (MW: 40,000; Polysciences). After 48 h, cells were harvested and sonicated in lysis buffer (50 mM HEPES [pH 8.0], 0.3 M NaCl, 0.2% NP40, protease inhibitor). After centrifugation, the ANTI-FLAG M2 Affinity Gel (A2220; Sigma-Aldrich) was added to the supernatant and incubated at 4 °C for 4 h. After washing the beads, SDS sample buffer was added, and the immunoprecipitated proteins were analyzed by western blotting.

Protein-protein interactions between Mkl1/2 and Foxo1/3/4 or glucocorticoid receptors were also measured using NanoBiT PPI Starter Systems (Promega). 293FT cells were seeded in 96-well white wall microplates and co-transfected with the LgBiT and SmBiT plasmids. The SmBiT-HaloTag plasmid was used for the background measurements. At 24 h post-transfection, luminescent substrate was added, and luminescence was measured using the GloMax Discover Microplate Reader (Promega).

### Gene expression analyses

For RNA extraction, mice were euthanized, and WAT or skeletal muscle was isolated. TRIzol reagent (Thermo Fisher Scientific) was then added. The tissues were homogenized using a Micro Smash MS-100R (TOMY), and the FavorPrep Blood/Cultured Total RNA Mini Kit (FAVORGEN) was used for RNA extraction from cultured cells. In both cases, RNA was extracted according to the manufacturer’s instructions. Purified RNA was analyzed using a microarray (3D-Gene; TORAY) or RT-qPCR. For RT-qPCR, purified RNA was reverse-transcribed using ReverTra Ace (TOYOBO). Gene expression was measured using the SsoAdvanced Universal SYBR Green Supermix (Bio-Rad) with primer sets specific for each cDNA. Most primers used for RT-qPCR were pre-designed using PrimerBank, but primers for 18S rRNA, Mkl2, and ChIP-qPCR were designed using Primer-BLAST (NIH). The primers used are listed in Supplementary Table 2.

The gene expression profiles of the gastrocnemius muscle in aging or cancer cachexia were re-analyzed using RNA-seq data (GEO accession numbers: GSE145480 and GSE65936) in the GEO database. For upstream analyses, an Ingenuity Pathway Analysis was used to analyze the list of genes with variable expression obtained from the microarray.

### Viral vector production

For adenovirus production, the adenovirus genome was excised from the cosmid vector using the restriction enzymes BspT104I or PacI, and the excised DNA was purified and transfected into Adeno-X 293 cells (TaKaRa). When late CPE was observed, the cells and culture medium were collected, and the virus was extracted by freezing and thawing six times. In addition, the cell infection and virus extraction cycle was repeated twice to produce a high-titer adenovirus solution.

The adeno-associated virus MyoAAV.2A was produced using the AAV-MAX Helper-Free AAV Production System Kit (Thermo Fisher Scientific). At 72 h post-transfection, AAV was extracted with citrate buffer (55 mM citric acid, 55 mM sodium citrate, and 800 mM NaCl). The extract was neutralized with HEPES (pH 8.0) and precipitated with PEG 8000. The pellet containing AAV was resuspended and treated with Benzonase (Merck) for nucleic acid digestion. For further purification, density gradient ultracentrifugation was performed using the OptiPrep. The purified AAV was replaced with buffer in PBS, and the viral genome (vg) content was determined by RT-qPCR.

### Artificial microRNA (miRNA)

Artificial miRNAs that repress the expression of *Mkl1* or *Mkl2* were incorporated into the pAxcwit2-TetOne-eGFP, and adenoviruses encoding artificial miRNAs were generated. An miRNA cluster consisting of three miRNAs is located in the 3′-UTR of eGFP.

The sequence of each miRNA was as follows:

*Mkl1* miRNA #1: 5*’*-CTGTTCAAAGGCGGCATGATCCCTGTTTTGGCCACTGACTGACAGGGATCACCGCCTTTGAACAG-3*’*;

*Mkl1* miRNA #2: 5*’*-CTGATTTCTCGCTGGCAGACTTGGGTTTTGGCCACTGACTGACCCAAGTCTCAGCGAGAAATCAG-3*’*;

*Mkl2* miRNA #1: 5*’*-CTGAGAACTGCGAGACAACGTTCTGTTTTGGCCACTGACTGACAGAACGTTCTCGCAGTTCTCAG-3*’*;

*Mkl2* miRNA #2: 5*’*-CTGAGACCTTTCTGACTGGCTGAGGTTTTGGCCACTGACTGACCTCAGCCACAGAAAGGTCTCAG-3*’*.

### Statistical analyses

Measurements for each experiment are presented as the mean ± standard deviation. Statistical significance was determined by Student’s *t*-test, Dunnett’s test, or Tukey’s test using the JMP statistical software program (JMP), with a p-value <0.05 considered significant.

## Author Contributions

A.K. designed and performed the experiments and analyzed the data. A.K., K.H., R.S. and T.O. discussed and interpreted the analyses. The manuscript was written by A.K and edited by all authors.

## Acknowledgments

We thank T.O., K.H., and R.S. laboratory members for technical assistance, comments on the manuscript, and scientific discussions. We thank Masafumi Inui for scientific discussions. We thank the Institute of Development, Aging and Cancer (IDAC), Tohoku University, and the Kobe Biomedical Research Foundation for providing the aged mice. This work was supported by JSPS Grant-in-Aid for Scientific Research 21K11386 (A.K.); The Ito Foundation (A.K.); Daiwa Securities Foundation (A.K.); CASIO Science Promotion Foundation (A.K.); TOBE MAKI Scholarship Foundation (A.K.); and AMED 19gm0810001 (T.O.).

---

**Extended Data Fig. 1:**
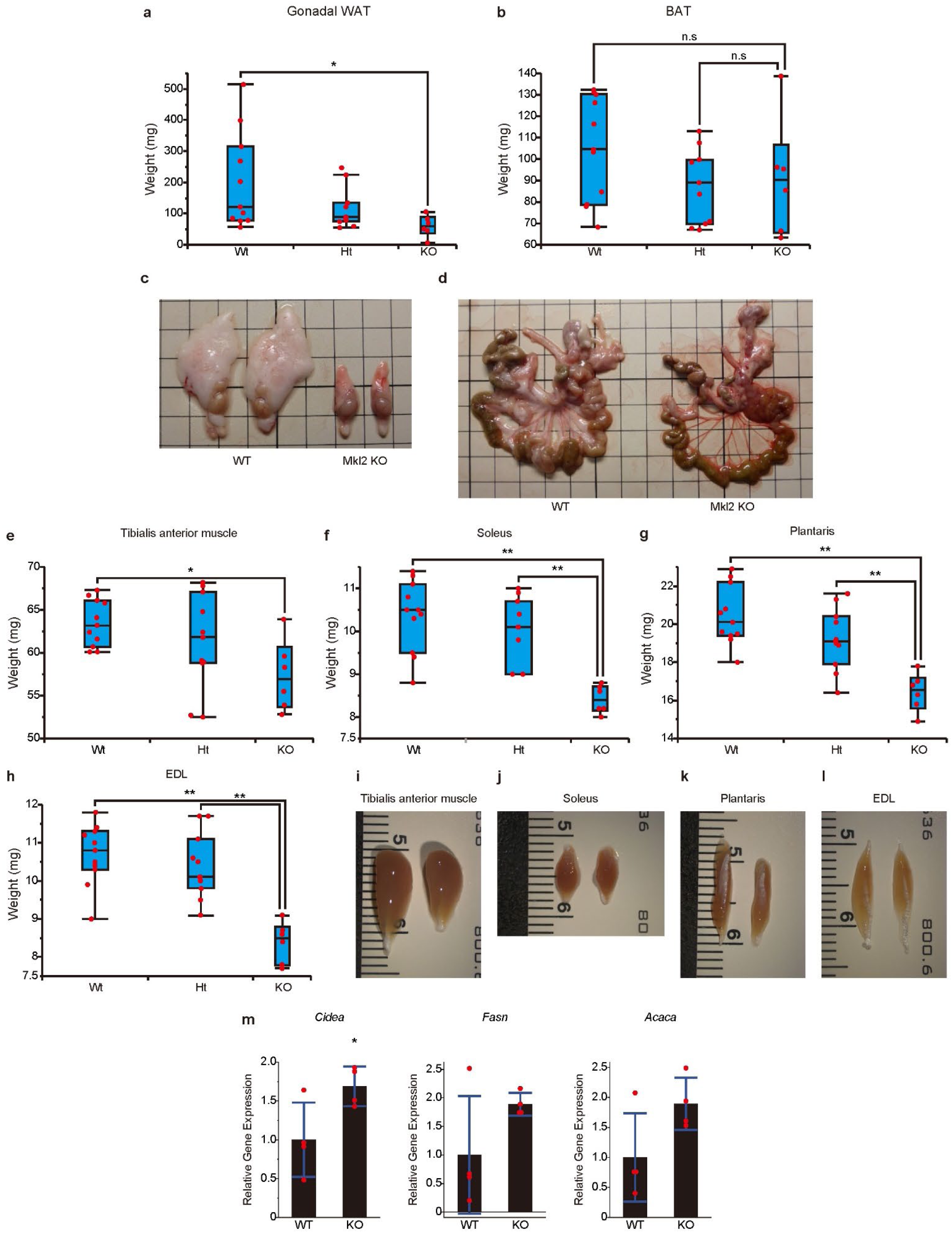
Both white adipose tissue and skeletal muscle mass are reduced in *Mkl2*-knockout mice. **a**, Gonadal WAT weight of *Mkl2* KO mice and their littermates. **b**, Brown adipose tissue (BAT) weight of *Mkl2* KO mice and their littermates. **c, d** WAT of *Mkl2* KO mice and their littermates. Epididymal WAT (**c**) and mesenteric fat (**d**). **e-h**, Skeletal muscle weight of *Mkl2* KO mice and their littermates. **i-l,** tibialis anterior (**i**), soleus (**j**), plantaris (**k**), and extensor digitorum longus muscles (**l**) of *Mkl2* KO mice and their littermates. **m**, Expression of Foxo1 target genes in the inguinal WAT of *Mkl2* KO mice and their littermates. Data are shown as the mean ± s.d. (n=4). **a, b**, **e-h**, Data are presented as box plots (21-22 weeks old, female mice; WT, n=11; Ht, n=11; KO, n=6). **a**, **b**, **e**-**h**, **m**, significance (*p <0.05 or **p <0.01) was calculated using Student’s *t*-test (**m**) or Dunnett’s multiple comparison test (**a**, **b**, **e**-**h**). n.s: not significant.

**Extended Data Fig. 2:**
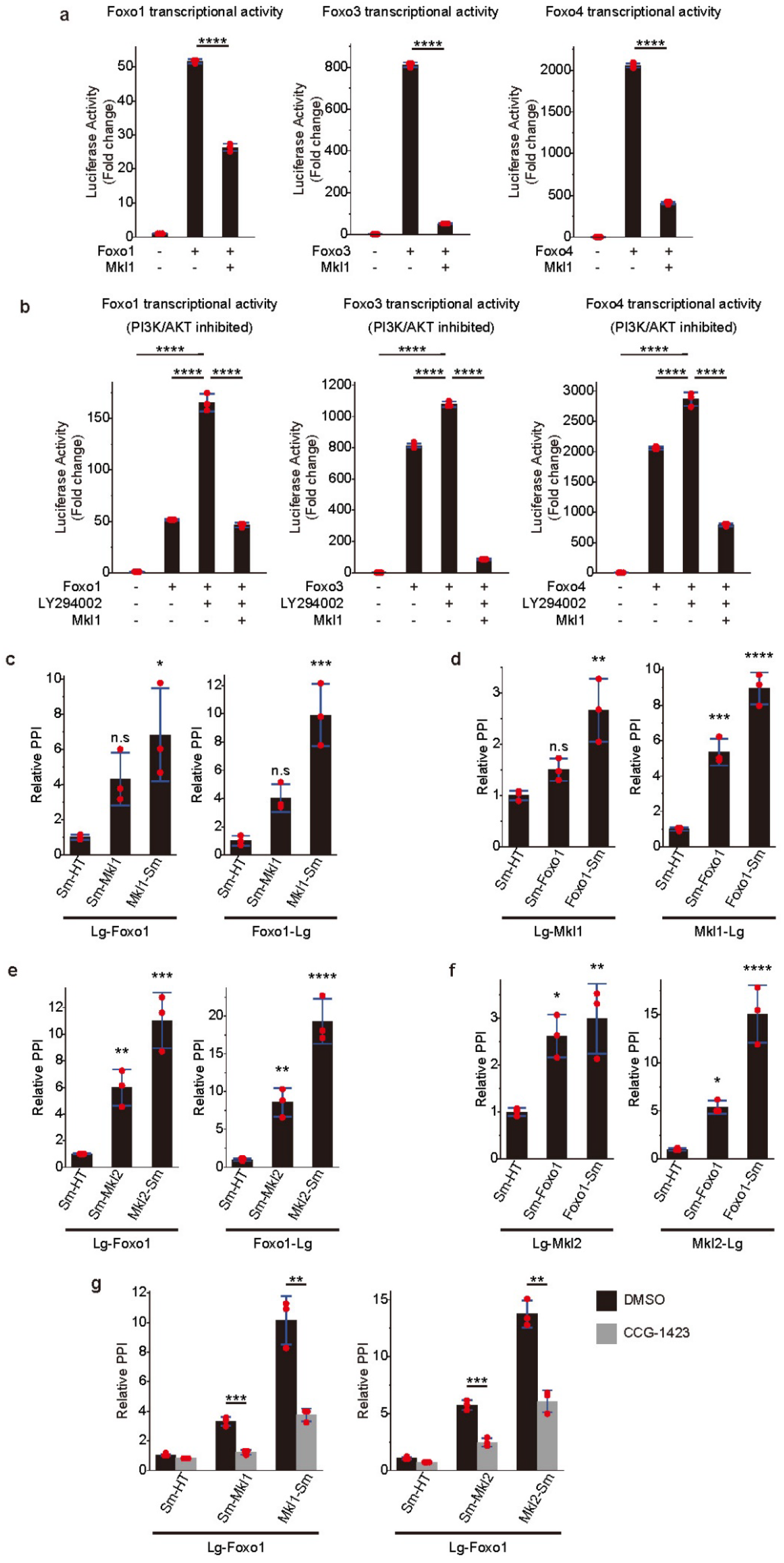
Mkl1 and Mkl2 bind directly to FoxO proteins and repress their transcriptional activity. **a, b**, Inhibitory effects of Mkl1 on the transcriptional activity of Foxo1, Foxo3, and Foxo4 using luciferase reporters with FoxO-responsive elements in the normal state (**a**) or with the PI3K/AKT pathway suppressed by LY294002 treatment (**b**). **c-f**, Protein-protein interaction (PPI) between Foxo1 and Mkl1 (**c, d**) or Mkl2 (**e, f**) detected by split luciferase (NanoBiT). PPI was measured for all possible combinations (Lg: LgBiT tag, Sm: SmBiT tag, and HT: Halotag). The relative PPI signal was calculated based on the signal between the LgBiT-tagged protein and the SmBiT-Halotag. **g** Changes in the interaction of Foxo1 with Mkl1 or Mkl2 by adding CCG-1423 were measured in different combinations, as shown in Fig. 2f. **a**-**g**, Data are shown as the mean ± s.d. of n = 3 cultures per condition. Significance (*p <0.05, **p <0.01, ***p <0.001, and ****p <0.0001) was calculated using Student’s *t*-test (**a**, **g**) or Dunnett’s multiple comparison test (**b-f**). n.s: not significant.

**Extended Data Fig. 3:**
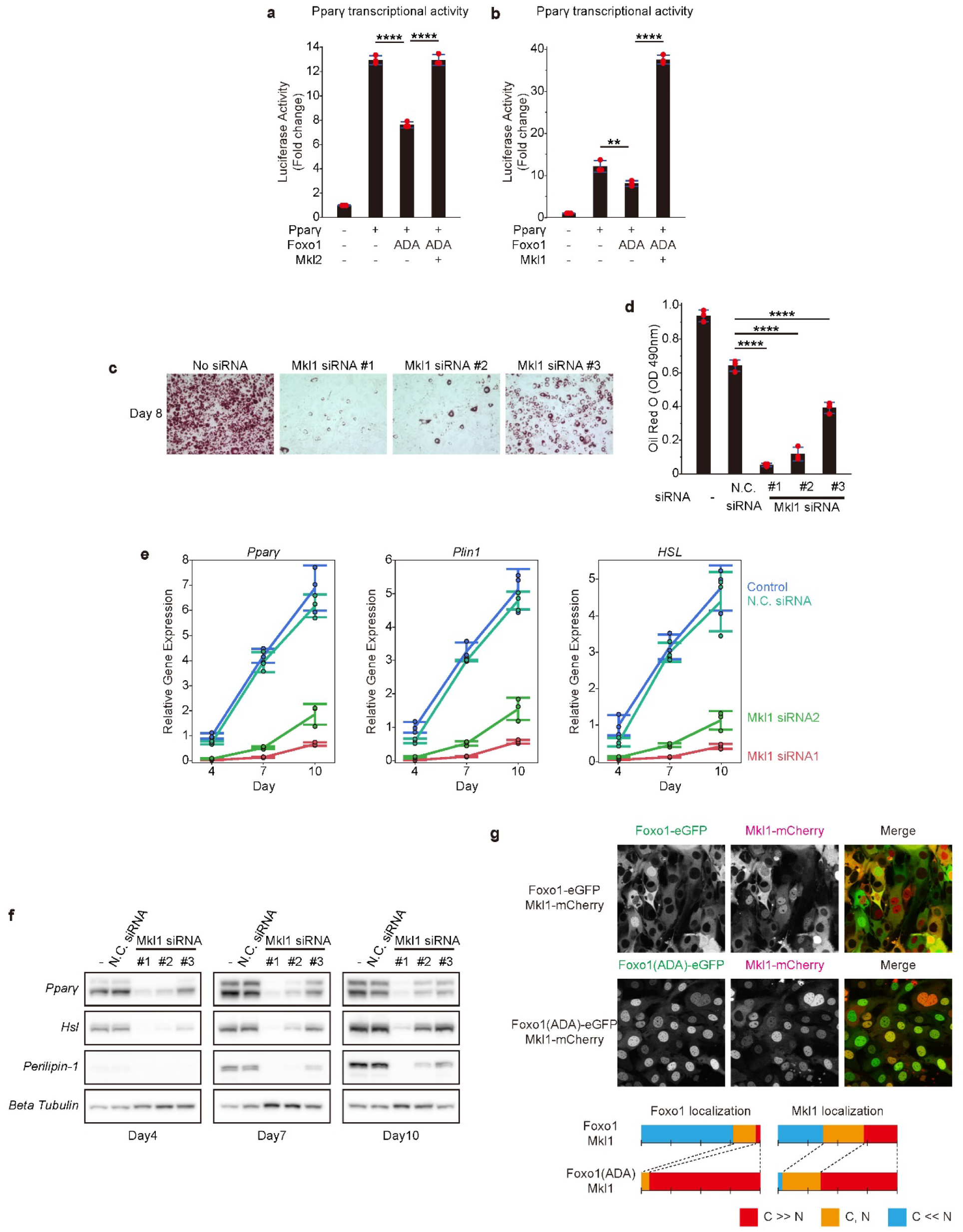
Knockdown of *Mkl1* impaired adipocyte differentiation. **a**, **b,** A luciferase assay using a reporter with *Pparγ*-responsive elements. Pparγ transcriptional activity was suppressed by a constitutively active mutant of Foxo1 (ADA) but restored by Mkl2 (**a**) and Mkl1 (**b**). **c**, **d**, Oil Red O staining of *Mkl1* knockdown 3T3-L1 cells at day 8 of differentiation (**c**). After staining, the dye was extracted with isopropanol and quantified by measuring the absorbance at 490 nm (**d**). **e**, mRNA expression of *Pparγ*, *Plin1* and *Hsl* on days 4, 7, and 10 after induction of differentiation of 3T3-L1 cells with or without *Mkl1* knockdown. mRNA expression is shown as the relative change from the average expression of the control on day 4. **f**, Protein expression of *Pparγ*, *Plin1* and *Hsl* on days 4, 7, and 10 after induction of differentiation of 3T3-L1 cells with or without *Mkl1* knockdown. **g**, Subcellular localization of Mkl1 with Foxo1 or Foxo1(ADA) expression. The percentage of cells showing nuclear and cytoplasmic localization of Foxo1 and Mkl1 are shown below (C: cytoplasmic localization, N: nuclear localization). **a, b, d**, Data are shown as the mean ± s.d. of n = 3 cultures per condition. Significance (**p <0.01 or ****p <0.0001) was calculated using Dunnett’s multiple comparison test.

**Extended Data Fig. 4:**
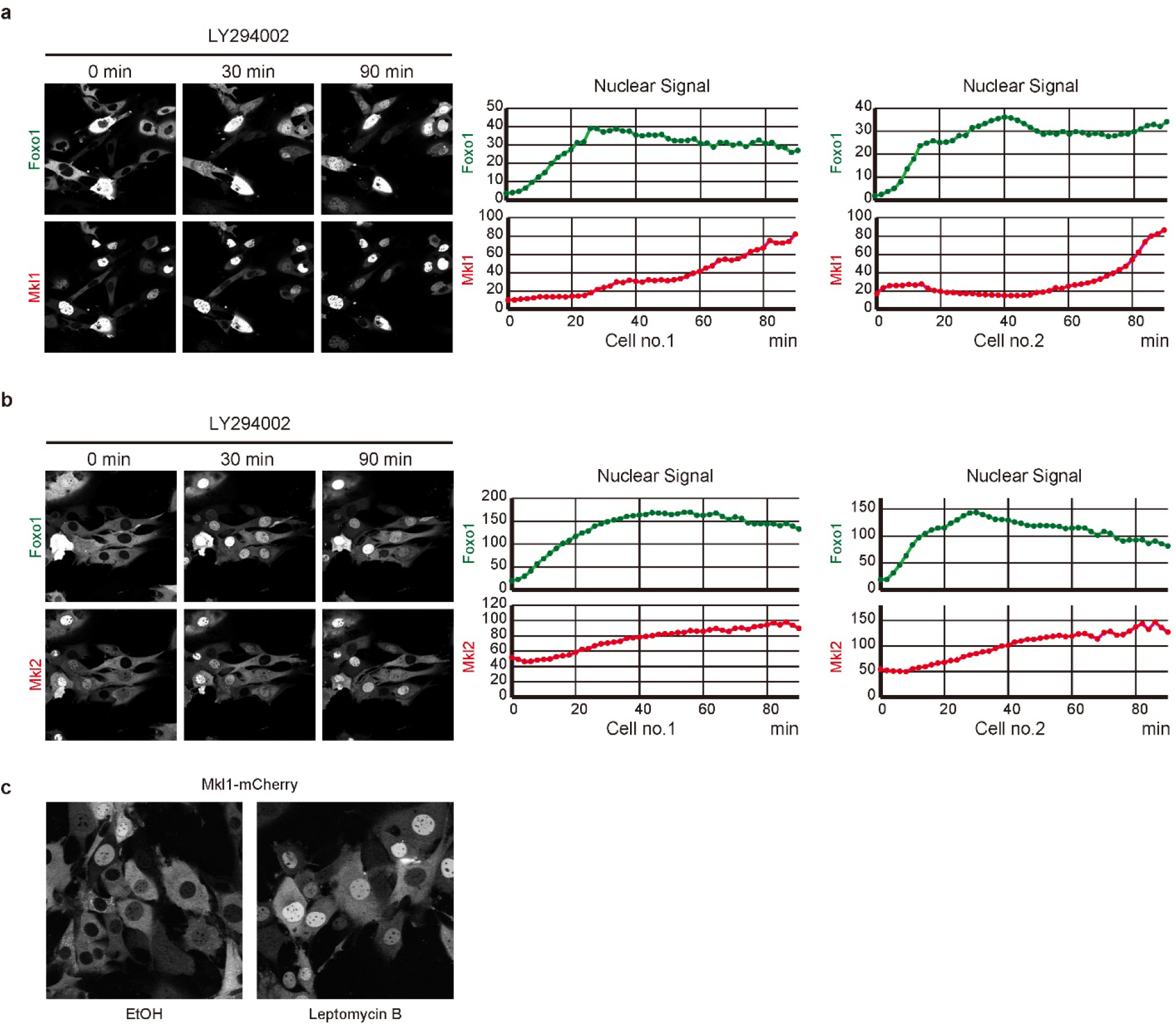
Mkl1 and Mkl2 accumulate in the nucleus in response to Foxo1 activation. **a**, **b**, Changes in the localization of Foxo1 and Mkl1 (**a**) or Mkl2 (**b**) after the addition of LY294002, a PI3K inhibitor. Changes in the nuclear signal over time are shown on the right for the two cells showing representative behavior. **c** Changes in Mkl1 localization after treatment with or without leptomycin B.

**Extended Data Fig. 5:**
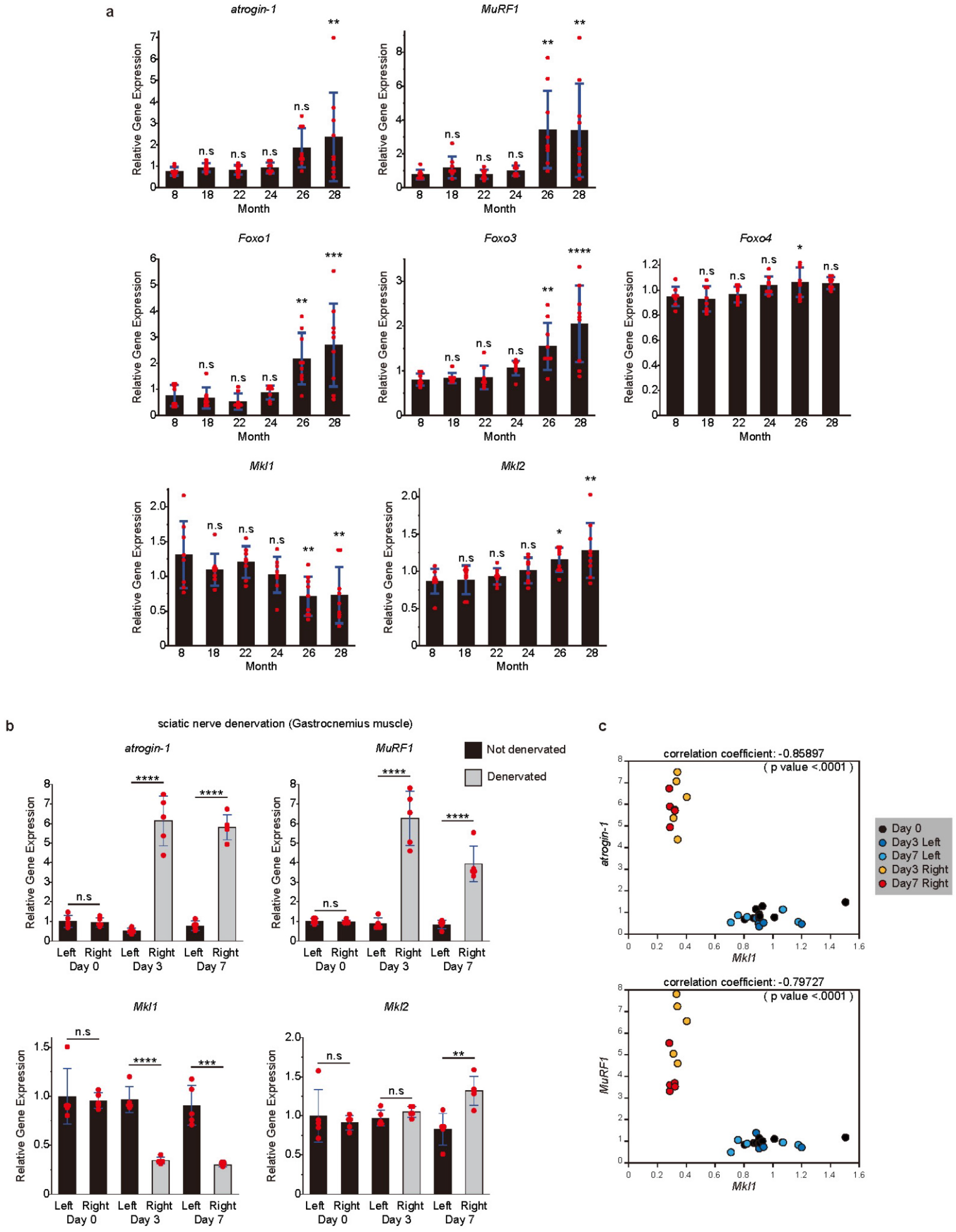
Expression changes of *atrogin-1*, *MuRF1*, *Foxo1/3/4*, and *Mkl1/2* in muscle atrophy induced by aging or sciatic nerve denervation. **a**, Expression profiles of *atrogin-1*, *MuRF1*, *Foxo1*, *Foxo3*, *Foxo4*, *Mkl1*, and *Mkl2* in gastrocnemius muscle of 8-, 18-, 22-, 24-, 26-, and 28-month old mice. These data were obtained from a re-analysis of the GEO dataset (GEO accession number: GSE145480). **b**, Changes in expression of *atrogin-1*, *MuRF1*, *Foxo1*, *Foxo3*, *Foxo4*, *Mkl1*, and *Mkl2* in the left or right gastrocnemius muscle of mice with sciatic nerve denervation of the right hindlimb. Day = number of days after sciatic nerve denervation. **c**, Inverse correlation between *Mkl1* and atrogene expression in the gastrocnemius muscle of mice with right hindlimb sciatic nerve denervation. Each point represents the relative gene expression in either the left or right mouse gastrocnemius muscle. Relative gene expression was calculated based on the average expression in the left leg before sciatic nerve denervation (Day 0). **a**, **b** Data are shown as the mean ± s.d. (n=5). Significance (*p <0.05, **p <0.01, ***p <0.001, or ****p <0.0001) was calculated using Student’s *t*-test (**b**) or Dunnett’s test (**a**) for multiple comparisons. n.s: not significant.

**Extended Data Fig. 6:**
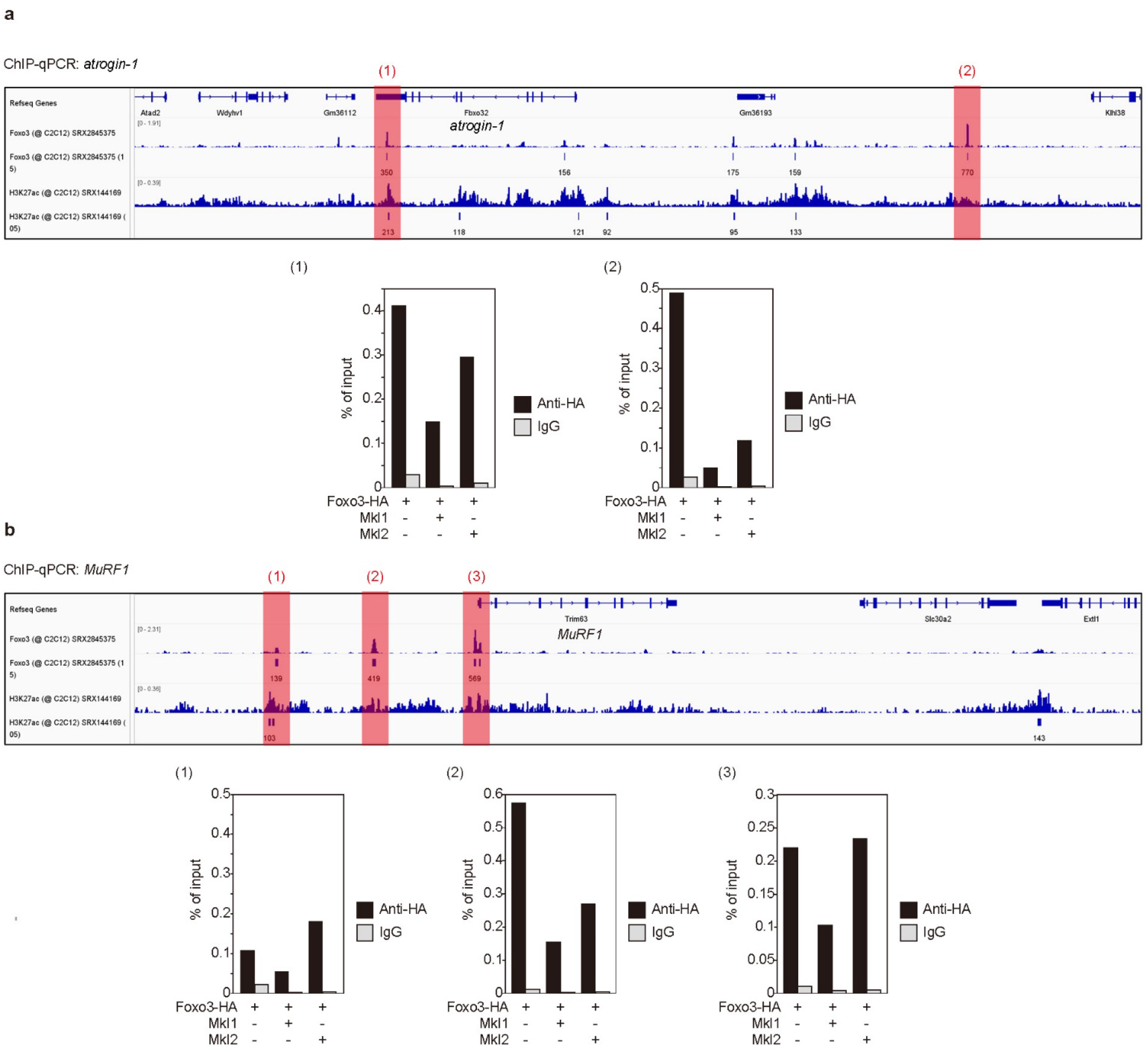
Foxo3 binding to the cis-regulatory region of atrogenes is inhibited by Mkl1 or Mkl2. **a**, Changes in Foxo3 binding to FoxO binding sites on genomic regions near *atrogin-1* (**a**) or *MuRF1* (**b**) due to *Mkl1* or *Mkl2* expression, as measured by chromatin immunoprecipitation (ChIP-qPCR).

**Extended Data Fig. 7:**
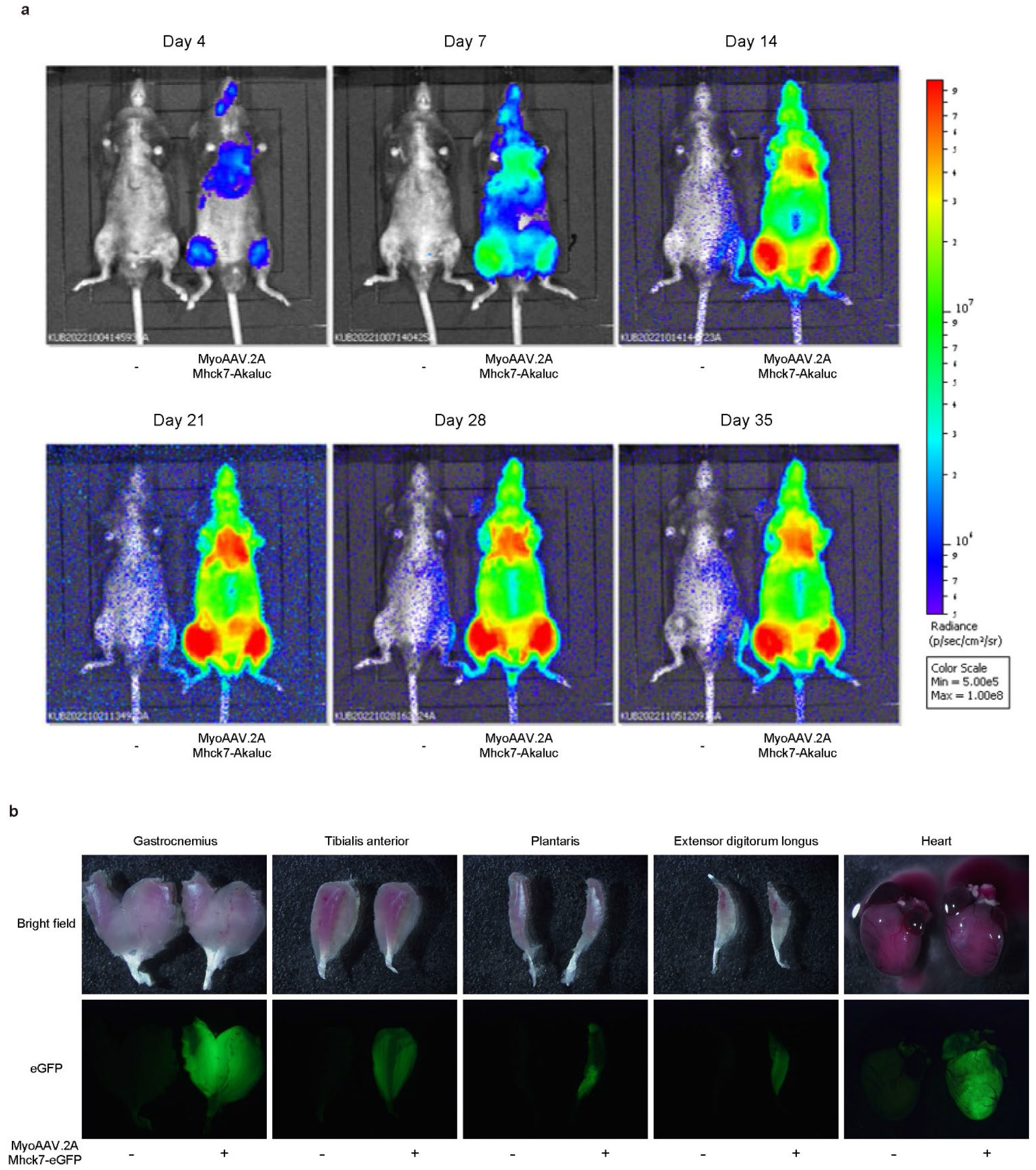
MyoAAV.2A efficiently delivers AkaLuc-eGFP to skeletal and cardiac muscle. **a**, Changes in AkaLuc expression over time in mice systemically treated with MyoAAV.2A (Mhck7-AkaLuc-eGFP). Bioluminescence was measured using an IVIS Lumina II imaging system. **b**, eGFP expression observed in each muscle and hearts of mice systemically treated with MyoAAV.2A (Mhck7-AkaLuc-eGFP).

**Extended Data Fig. 8:**
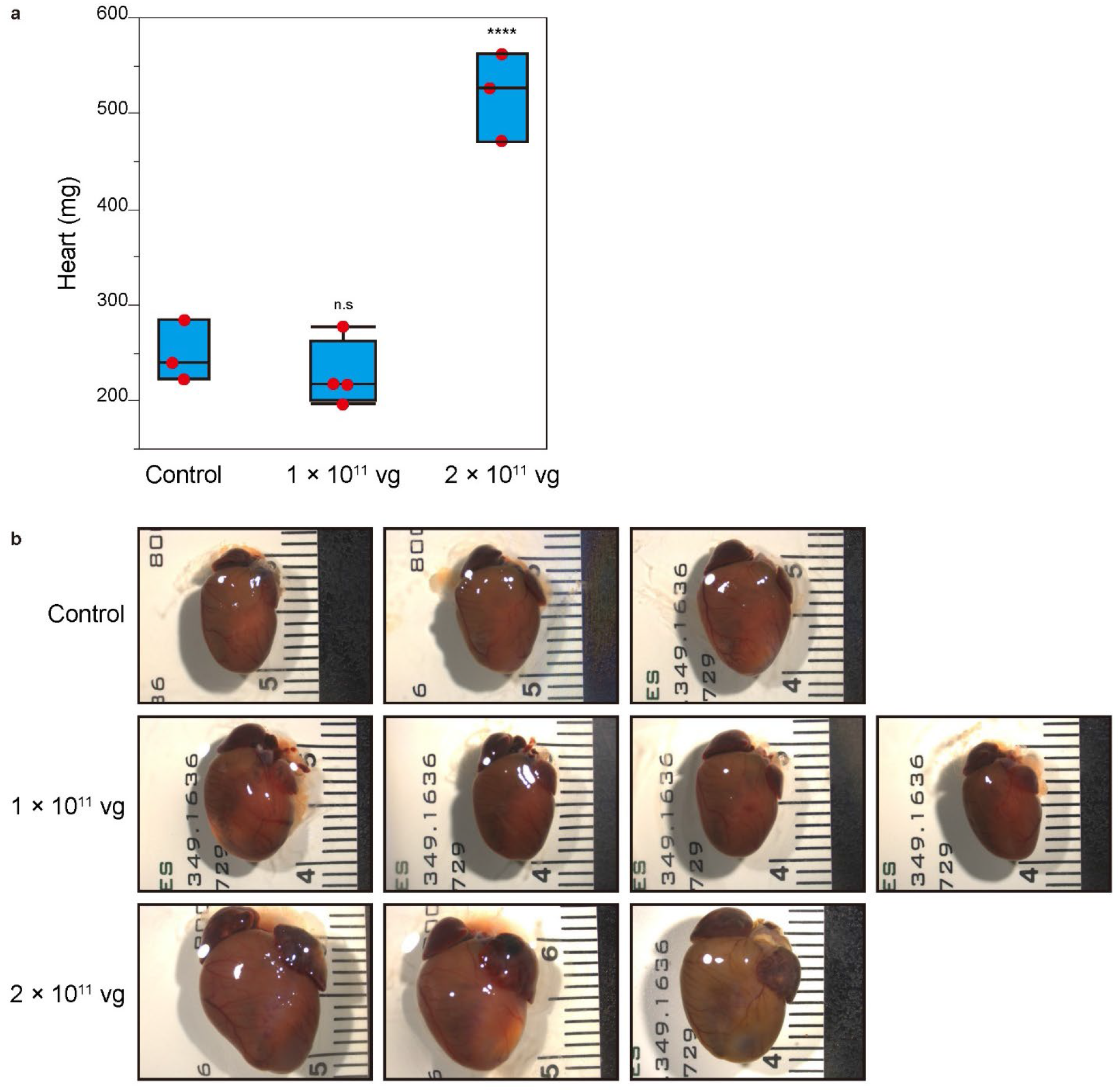
Cardiac hypertrophy was induced by MyoAAV-mediated expression of *Mkl1* in the myocardium. **a, b**, Heart weight (**a**) and morphology (**b**) of mice systemically treated with MyoAAV.2A (Mhck7-*Mkl1*) at different viral concentrations. Data are shown as box plots (n=3-4). Significance (****p <0.0001) was calculated using Dunnett’s test for multiple comparisons. n.s: not significant.

**Extended Data Fig. 9:**
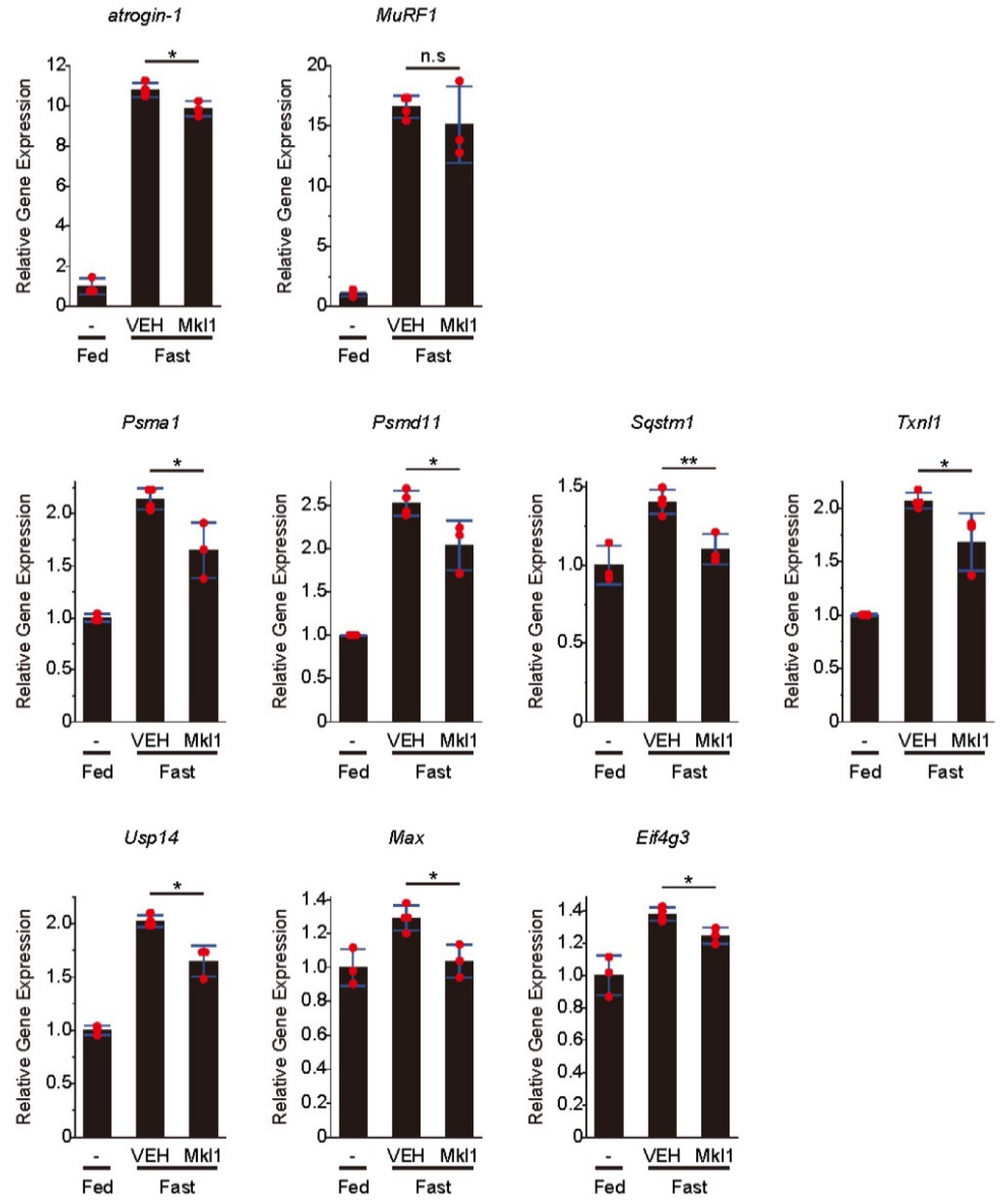
Supplementation of *Mkl1* suppresses the expression of atrogenes induced by fasting. A comparison of atrogene expression in the gastrocnemius muscle of mice expressing *Mkl1* by MyoAAV.2A and fasted for 24 h. Data are shown as the mean ± s.d. (n=3-4). Significance (*p <0.05 or **p <0.01) was calculated using Student’s *t*-test. n.s: not significant.

**Supplementary Movie 1**

Time-lapse imaging of Foxo1-eGFP and Mkl2-mCherry localization in 3T3-L1 preadipocytes after addition of the PI3K inhibitor wortmannin. The green channel (left) represents the Foxo1-eGFP expression, and the red channel (center) represents Mkl2-mCherry. The merged image is shown on the right side. This video is shown in Fig. 3h.

**Supplementary Movie 2**

The nuclear accumulation of Mkl2-eGFP during contraction of C2C12-derived myotubes 3 h after initiation of electrical pulse stimulation (EPS). Adenovirus was used to express Mkl2-eGFP in myotubes, and EPS was applied at a frequency of 1 Hz. This movie corresponds to the figures on the right side of Figure 8a.

